# Variant Phasing and Haplotypic Expression from Single-molecule Long-read Sequencing in Maize

**DOI:** 10.1101/654533

**Authors:** Bo Wang, Elizabeth Tseng, Primo Baybayan, Kevin Eng, Michael Regulski, Yinping Jiao, Liya Wang, Andrew Olson, Kapeel Chougule, Peter Van Buren, Doreen Ware

**Author notes:** Author for correspondence: Doreen Ware.

## Abstract

Haplotype phasing of genetic variants in maize is important for interpretation of the genome, population genetic analysis and functional genomic analysis of allelic activity. Accordingly, accurate methods for phasing the full-length isoforms are essential for functional genomics studies. We performed an isoform-level phasing study in maize, using two inbred lines and their reciprocal crosses, based on the single-molecule full-length cDNA sequencing. To phase and analyze the full-length transcripts between hybrids and parents, we developed a tool called IsoPhase. Using this tool, we validated the majority of SNPs called against matching short-read data and identified cases of allele-specific, gene-level and isoform-level expression. Our results revealed that maize parental lines and hybrid lines exhibit different splicing activities. After phasing 6,907 genes in two reciprocal hybrids using embryo, endosperm and root tissues, we annotated the SNPs and identified large-effect genes. In addition, based on single-molecule sequencing, we identified parent-of-origin isoforms in maize hybrids, distinct novel isoforms in maize parent and hybrid lines, and imprinted genes from different tissues. Finally, we characterized variation in cis- and trans-regulatory effects. Our study provides measures of haplotypic expression that could increase accuracy in studies of allelic expression.

## Introduction

Phasing of genetic variants is crucial for identifying putative causal variants and characterizing the relationship between genetic variation and phenotype^1^. Maize is a diploid organism that not only has high genetic diversity, but also exhibits allele-specific expression (ASE)^2,3,4^, i.e., unequal transcription of parental alleles. The effects of ASE vary by cell and tissue type, developmental stage, and growth conditions^5^. Because alleles from the same gene can generate heterozygous transcripts with distinct sequences, a comprehensive analysis of ASE is necessary in order to achieve a thorough understanding of transcriptome profiles^6, 7^. Previously, ASEs were studied using short-read RNA-seq, which could quantify alleles at the SNP level but was unable to provide full-length haplotype information^8, 9^. Third-generation sequencing technologies such as PacBio and Oxford Nanopore offer full-length transcript sequencing that eliminates the need for transcript reconstruction, and these methods have been widely adopted for genome annotations^10,11,12,13,14^. However, only a handful of studies have used long reads for isoform-level haplotyping^6^.

In this study, we used the Pacific Biosciences Sequel platform to produce a single-molecule full-length cDNA dataset for two maize parental lines, the temperate line B73 and the tropical line Ki11, as well as their reciprocal hybrid lines (B73 × Ki11; Ki11 × B73). We also sequenced RNAs from the same tissues using 150-bp paired-end (PE) sequencing on the Illumina platform. In addition, we developed a tool called IsoPhase to phase the allelic isoforms in the hybrids based on the single-molecule transcriptome datasets. This is the first full-length isoform phasing study in maize, or in any plant, and thus provides important information for haplotype phasing in other species, including polyploid species. Using IsoPhase, we demonstrated that haplotype phasing by full-length transcript sequencing can reveal allele-specific expression in maize reciprocal hybrids. Our approach does not require parental information (although parental data could be used to assign maternal and paternal alleles) and can be applied to exclusively long-read data. Moreover, we showed that single molecules could be attributed to the alleles from which they were transcribed, yielding accurate allelic-specific transcriptomes. This technique allows the assessment of biased allelic expression and isoform expression.

## Results

### Full-Length Transcript Sequencing and Bioinformatics Pipeline

Reciprocal hybrids of the maize inbred lines B73 and Ki11 exhibited dramatic heterosis in plant height, primary and lateral root number, biomass and 100-kernel Weight (Supplementary Fig. 1a-h), making this group well suited to the allelic study of heterosis. To qualitatively assess the gene expression profiles of various tissues, we extracted high-quality RNA from the root, embryo, and endosperm of B73, Ki11, and their reciprocal crosses (B73 × Ki11, Ki11 × B73), and then subjected the RNA to reverse transcription. Tissue-specific barcodes were added before pooling for subsequent amplification. Barcoded SMRTBell libraries were sequenced on the PacBio Sequel 1M platform with 15 SMRT Cells using 2.1 chemistry, yielding 4,898,979 circular consensus sequences (CCS, also called reads of insert). We pooled the reads from all four lines and processed them using the IsoSeq3 workflow (Fig. 1); 76.3% (3,739,812) of reads were classified as full-length insert transcripts based on the presence of barcoded primers and poly(A) tails (Supplementary Table 1 and Supplementary Table 2). IsoSeq3 processing yielded 250,168 full-length, high-quality (HQ) consensus transcript sequences.

**Figure 1:**
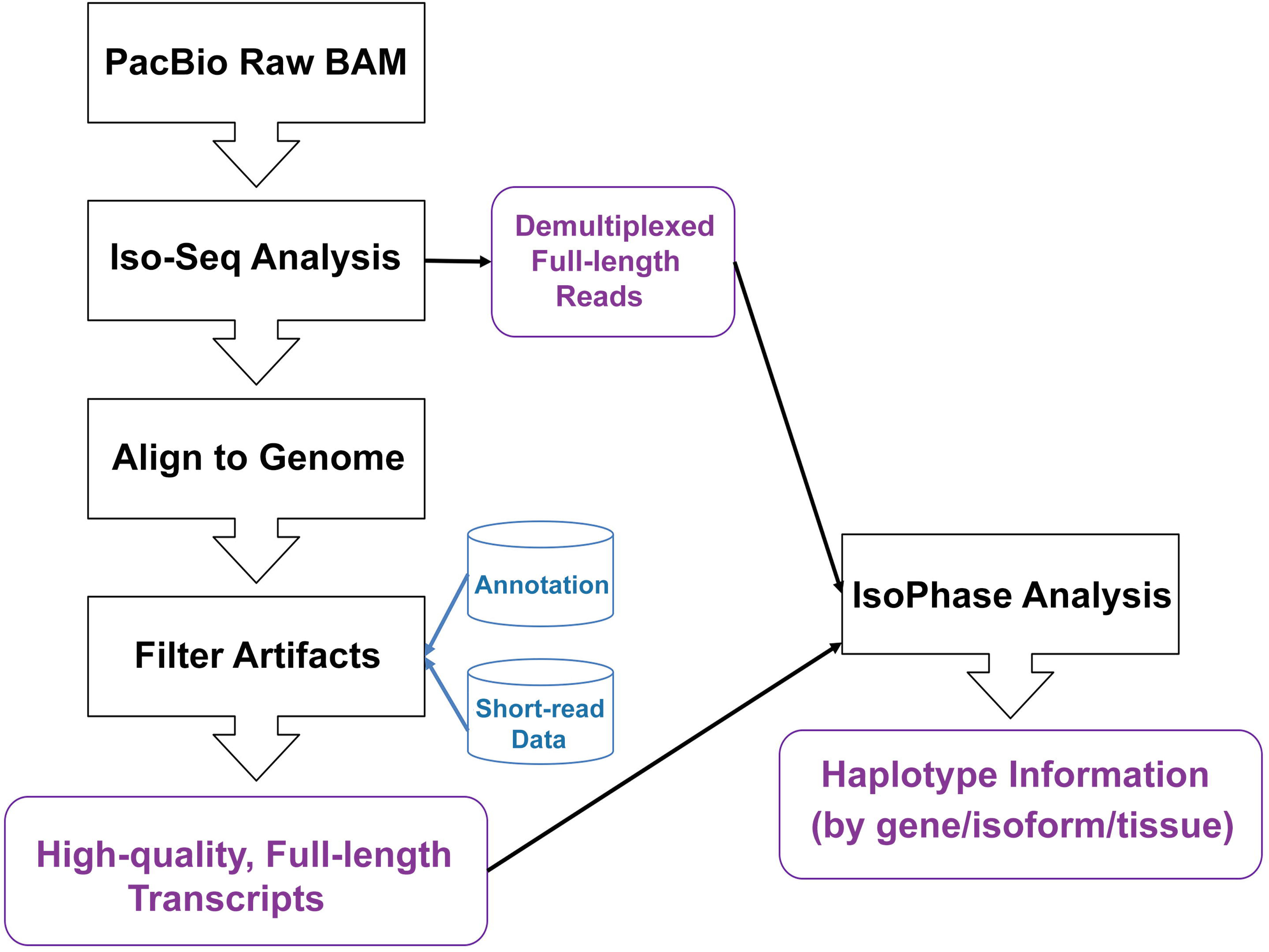
Full-length Transcript Sequencing Workflow. Full-length transcript data are generated on the PacBio Sequel platform following the recommended procedure for Iso-Seq multiplexed library preparation. Iso-Seq analysis demultiplexes each read and removes the barcodes, cDNA primers, and poly(A) tail. The demultiplexed reads are pooled and run through an isoform-level clustering algorithm that produces high-quality transcript sequences, which is then aligned to the reference genome and removed of artifacts. The genome-mapped genes are then used to run the IsoPhase analysis to call SNPs and haplotypes.

The HQ transcript sequences were then mapped to the maize RefGen_v4 genome assembly^14^ using minimap2^15^. Of the 250,168 HQ sequences, 248,424 (99.3%) were mapped to the genome, among which 229,757 (91.8%) were selected according to two criteria: min-coverage 99% and min-identity 95%. These sequences were further collapsed into 90,419 non-redundant transcripts (Supplementary Table 3). Using the QC tool SQANTI^15^, we discarded 15,301 (16.9%) of the non-redundant transcripts due to intra-priming, RT switching, or non-canonical junctions unsupported by matching short-read data (Supplementary Table 4). Genome-wide BLASTN to the NCBI RefSeq NR database revealed that 523 (30%) of the 1,744 unmapped sequences fell into gaps in the assembly, whereas the remaining sequences could be mapped to other organisms, implying that they represented biological contamination from endophytes or other sources (Supplementary Table 5). Our final dataset consists of 75,118 transcripts covering 23,412 gene loci, with lengths ranging from 80 to 11,495 bp and an average transcript length of 2,492 bp (Supplementary Fig. 2).

### Isoform Characterization in Maize B73, Ki11, and Reciprocal Lines

We used SQANTI to compare reference transcripts against the maize B73 RefGen_v4 annotation^14^, matching 20,068 of the 23,412 loci to a reference gene locus; the remaining 3,344 transcripts, with an average gene length of 2,350 bp, were deemed novel, which originated from B73 genome but missing in current annotation. Among these novel transcripts, 3,193 (95.5%) are intergenic, and 319 (4.5%) are antisense.

SQANTI classified the transcripts into six groups (Fig. 2a): a] 36,005 (47.9%) isoforms were FSM (Full Splice Match); b] 8,910 (11.9%) isoforms were ISM (Incomplete Splice Match); c] 13,521 (18.0%) were NIC (Novel In Catalog); d] 13,170 (17.5%) were NNC (Novel Not In Catalog); e] 319 (0.4%) were Antisense; and f] 3,193 (4.3%) were Intergenic. The Iso-Seq data can recover full-length reference transcripts up to 10kb in length (Fig. 2b), the shortest transcripts (< 1 kb) and novel genes were enriched for multi-exon transcripts (Fig. 2c-d), of which 44.3% (4,702) were non-coding. In addition, we identified alternative start and end sites for full splice matched transcripts (Fig. 2e-f).

**Figure 2:**
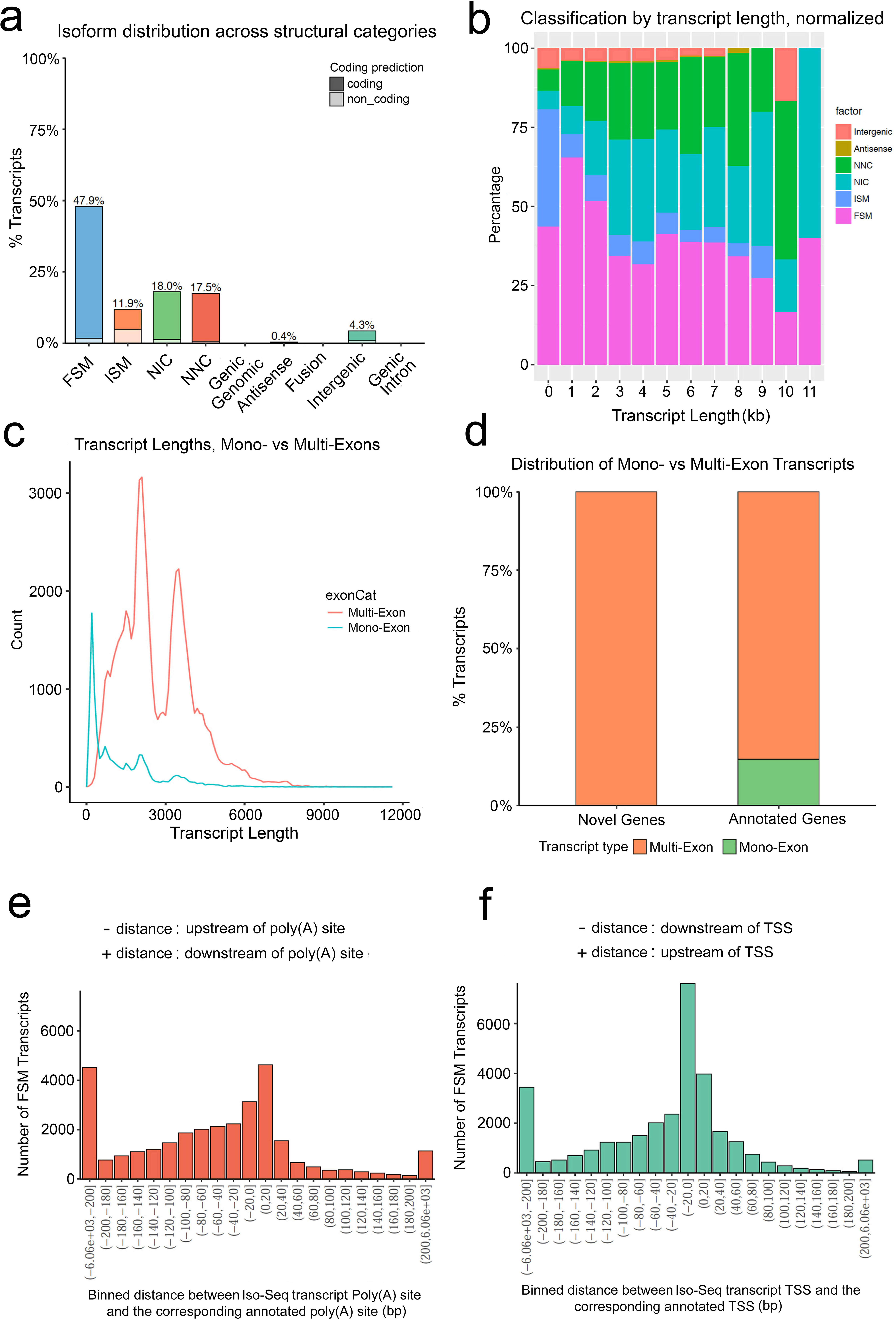
Iso-Seq transcript categorization against maize B73 RefGen_v4 annotation using the SQANTI software. a) Isoform distribution across structural categories. FSM=Full Splice Match: matches a reference transcript exon by exon. ISM=Incomplete Splice Match: matches a reference transcript exon by exon, but is missing one or more 5’ exons. NIC=Novel In Catalog: novel isoform using known splice sites. NNC=Novel Not In Catalog: novel isoform using at least one novel splice site. Because this analysis is performed after SQANTI filtering, all junctions must be in the annotation, canonical, or supported by matching short-read data. b) Classification by transcript length, normalized. c) Transcript lengths, mono vs multi-exon. d) Distribution of mono- vs multi-exon transcripts. Frequency distribution of FSM transcripts as a function of the binned distance between Iso-Seq transcript Poly(A) site and the corresponding annotated poly(A) site (**e**), and the binned distance between Iso-Seq transcription start site (TSS) and the corresponding annotated TSS (**f**).

We then demultiplexed the pooled transcripts by looking at the number of reads associated with each strain-tissue pair (ex: B73-root) for each transcript. If a sample had at least one full-length read associated with a transcript, it was considered to be expressed. Each sample contained between 20,000 and 30,000 expressed transcripts (Supplementary Table 6). To determine the degree of saturation of the data, we subsampled the full-length reads associated with the transcripts by strain and by tissue; the results of this analysis revealed that the Iso-Seq data were saturated at the gene level, but were still revealing additional diversity at the transcript level (Supplementary Fig. 3a-d). Note, however, that this saturation analysis was limited by the input library size and subject to sequencing bias on the PacBio platform.

Comparison of genes and isoforms between parents and hybrids revealed no significant patterns between the two parental lines and their reciprocal F1s (Supplementary Fig. 4). We identified a number of shared and genotype-specific genes and isoforms between the parental line and two hybrids, which were confirmed by short-reads data from two biological replicates based on Fleiss’ kappa score^17^ (Supplementary Fig. 5 and Supplementary Table 7). The results showed good agreement in classification between long and short reads. Investigation of the main splicing patterns revealed that intron retention was the predominant pattern across these four genotypes, and alternative last exon the least common (Supplementary Fig. 6a-c). Quantification using Illumina short reads revealed that most genes exhibited additive expression patterns in the three tissues. We also observed differences between the two reciprocal hybrids: specifically, Ki11 × B73 had more non-additive expression genes than B73 × Ki11 in root and endosperm tissues, with the exception in embryo (Supplementary Fig. 7a-c), which could contribute to phenotypic differences between these two hybrid lines. However, there were no significant differences in the number of isoforms between genes with additive expression and those with non-additive expression, nor were there differences in genes with non-additive expression between the two reciprocal hybrids (Supplementary Fig. 8a-c).

### Full-Length Transcripts Enable Accurate Haplotyping

To phase Iso-Seq transcripts, we developed a new tool called IsoPhase. For each gene, we aligned all full-length reads to the gene region, and then called SNPs individually (currently, IsoPhase only calls substitution SNPs). We then used the full-length read information to reconstruct the haplotypes and used a simple error correction scheme to obtain the two alleles (Fig. 3a). To determine which allele belongs to B73 or Ki11, we took advantage of the fact that all B73 reads must only express one allele, whereas all Ki11 reads must only express the other. Once the parental alleles were identified, we obtained the allelic counts for the F1 hybrids (Fig. 3b). We applied IsoPhase to the 9,463 genes that had at least 40 FL read coverage, of which 6,907 had at least one SNP and could be readily classified as the B73 or Ki11 alleles (Supplementary Data 1).

**Figure 3:**
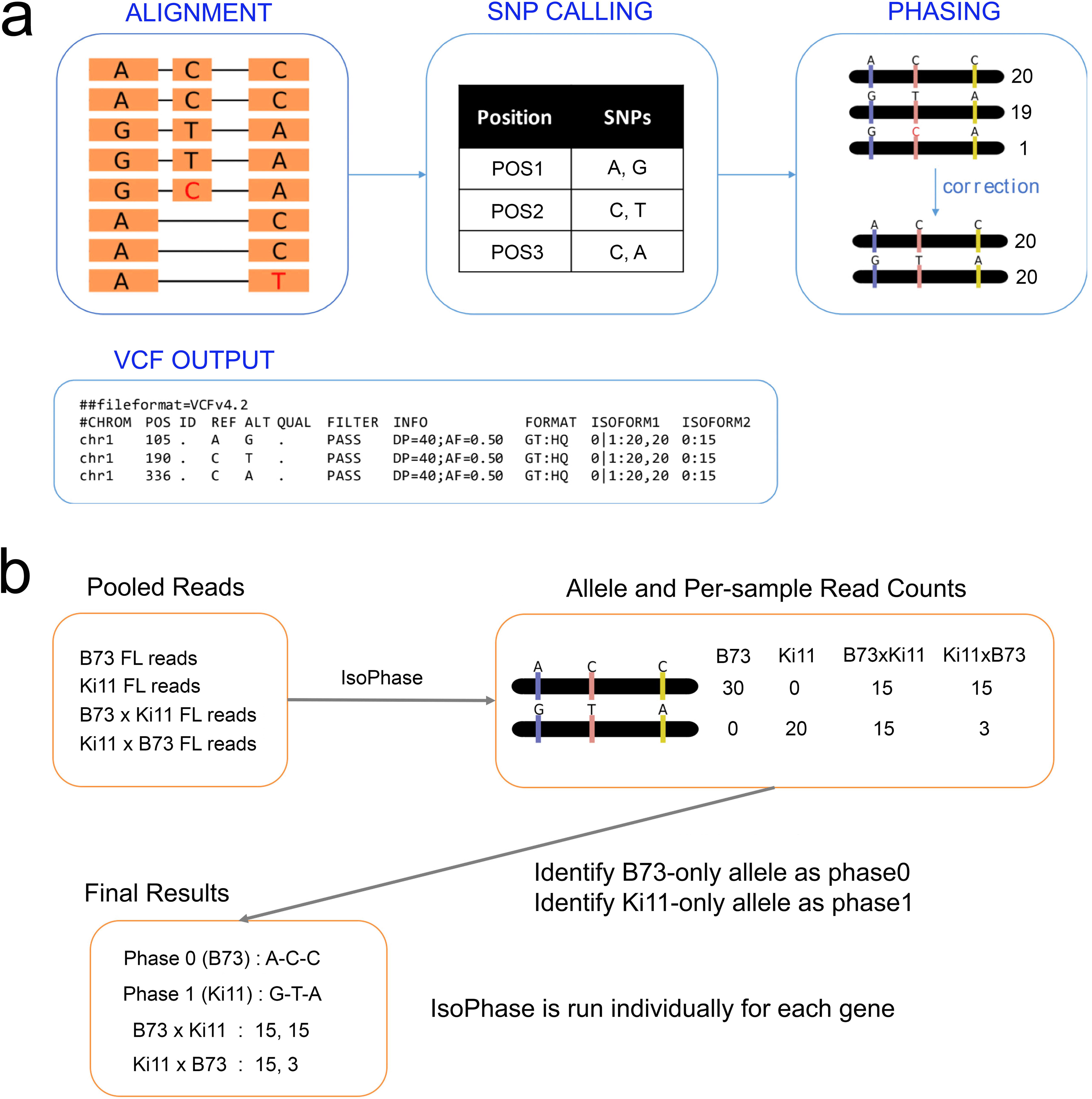
IsoPhase workflow. a) For each gene, full-length reads from all 12 samples are aligned to a gene region. SNPs are called individually for each position using Fisher’s exact test with the Bonferroni correction, applied with a p-value cutoff of 0.01. Only substitution SNPs are called. The full-length reads are then used to reconstruct the haplotypes, and a simple error-correction scheme is applied to obtain the two alleles. b) To determine which allele is derived from B73 vs. Ki11, we use the FL count information associate with the homozygous parents: B73 would only express the B73 allele, whereas Ki11 would only express the Ki11 allele.

We validated the SNPs called from IsoPhase using short-read data. Considering only substitution SNPs at positions for which there was at least 40 FL read coverage, 96% (74,280 of 77,540) of IsoPhase SNPs were validated by short-read data. The remaining 4% (3,260) of SNPs that were PacBio-specific were mostly due to insufficient coverage of the UTR regions by short-read data. Conversely, short reads identified an additional 26,774 SNPs in the regions that were not confirmed by IsoPhase. There were several reasons for this: (1) low or dropped coverage of Iso-Seq data; (2) alignment artifacts; and (3) indels masquerading as a series of consecutive SNPs. Both short read and Iso-Seq data showed variable coverage at the 5’ ends. In some cases, short-read data called additional 5’ SNPs (Fig. 4a). At positions where short reads called a SNP but Iso-Seq had sufficient coverage, however, read mis-alignment was a common issue (Supplementary Fig. 9a-b). In summary, we have confidence in the joint SNP calls, and attribute the unique SNP calls to either false negatives (due to low coverage) or false positives (due to mis-alignment).

**Figure 4:**
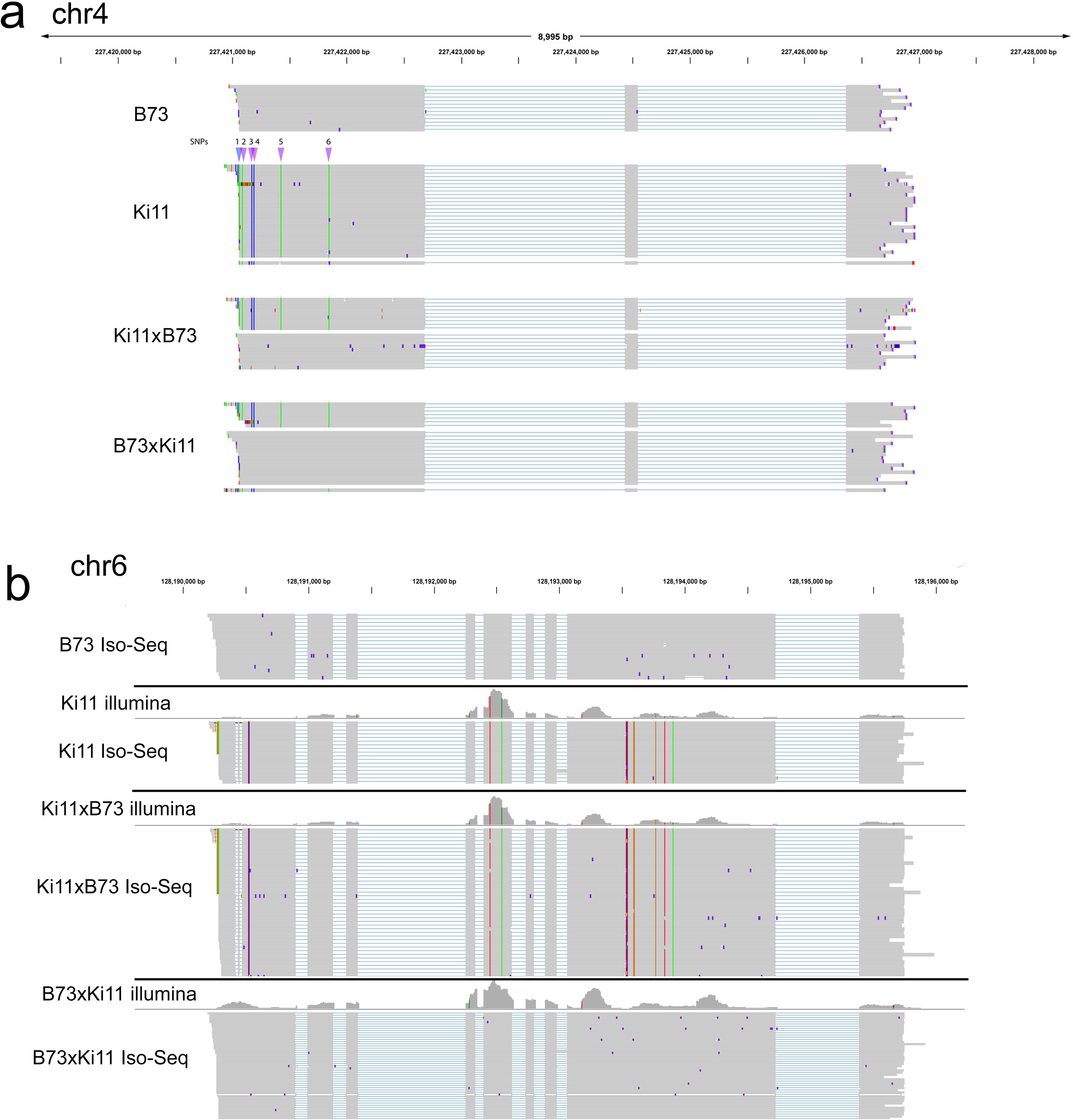
Example of phasing using IsoPhase. a) The gene PB.12426 (Zm00001d053356) phased by IsoPhase. The top two tracks show the B73 and Ki11 FL reads, and the bottom two tracks show the FL reads from the two F1 hybrids, with reads segregated by parental origin. SNPs are depicted between the B73 and Ki11 track, six SNPs are shown. SNPs #2–6 were called based on both long- and short-read data (purple); SNP #1 was missed by the long-read data due to reduced coverage (blue). b) IsoPhase can identify maternal-specific gene expression. The gene PB.16588 (Zm00001d037529) is expressed in both B73 and Ki11. However, in the F1 hybrids, only the maternal allele is expressed. In Ki11 × B73, the maternal allele is Ki11. In B73 × Ki11, the maternal allele is B73. Short-read data confirmed this maternal-specific gene expression.

### Full-Length Transcript Reads Reveal Allelic Specific Expression

An advantage of full-length transcript sequencing is the ability to characterize isoforms with haplotype information. Among the highly expressed genes, we observed cases of allelic-specific expression. For example, only the maternal allele of the gene Zm00001d037529 (PB.16588 gene from PacBio sequencing in this study) was expressed in the F1 hybrids, and both Iso-Seq and short-read data supported maternal-only expression (Fig. 4b). The Zm00001d040612 (PB.8517) gene provides a remarkable example of allelic-specific isoform expression. Its two most abundant isoforms were PB.8517.1 and PB.8517.4, which differ only in the last exon: PB.8517.4 had a single 3’ exon, whereas in PB.8517.1 the last exon is spliced into two exons. In B73, PB.8517.4 was expressed but PB.8517.1 was not, whereas in Ki11 the opposite pattern was observed. In both F1 hybrids, both isoforms were expressed, but only the Ki11 allele was detected for the PB.8517.1 isoform, whereas only the B73 allele was detected for the PB.8517.4 isoform (Fig. 5). Short-read junction data supported this observation. In total, we identified 221 monoallelic genes in embryo, 527 in endosperm, and 271 in roots (Supplementary Fig. 10a). Comparison of the number of isoforms in these monoallelic genes revealed differences among the three tissues: in embryo and root, the two reciprocal hybrids were inclined to exhibit the maternal effect pattern, whereas in endosperm, reciprocal hybrid B73 × Ki11 had more isoforms (Supplementary Fig. 10b). The isoform-level agreement in embryo, endosperm, and root tissue had kappa scores of 0.979, 0.965, and 0.973, respectively, indicating good agreement between long and short reads.

**Figure 5:**
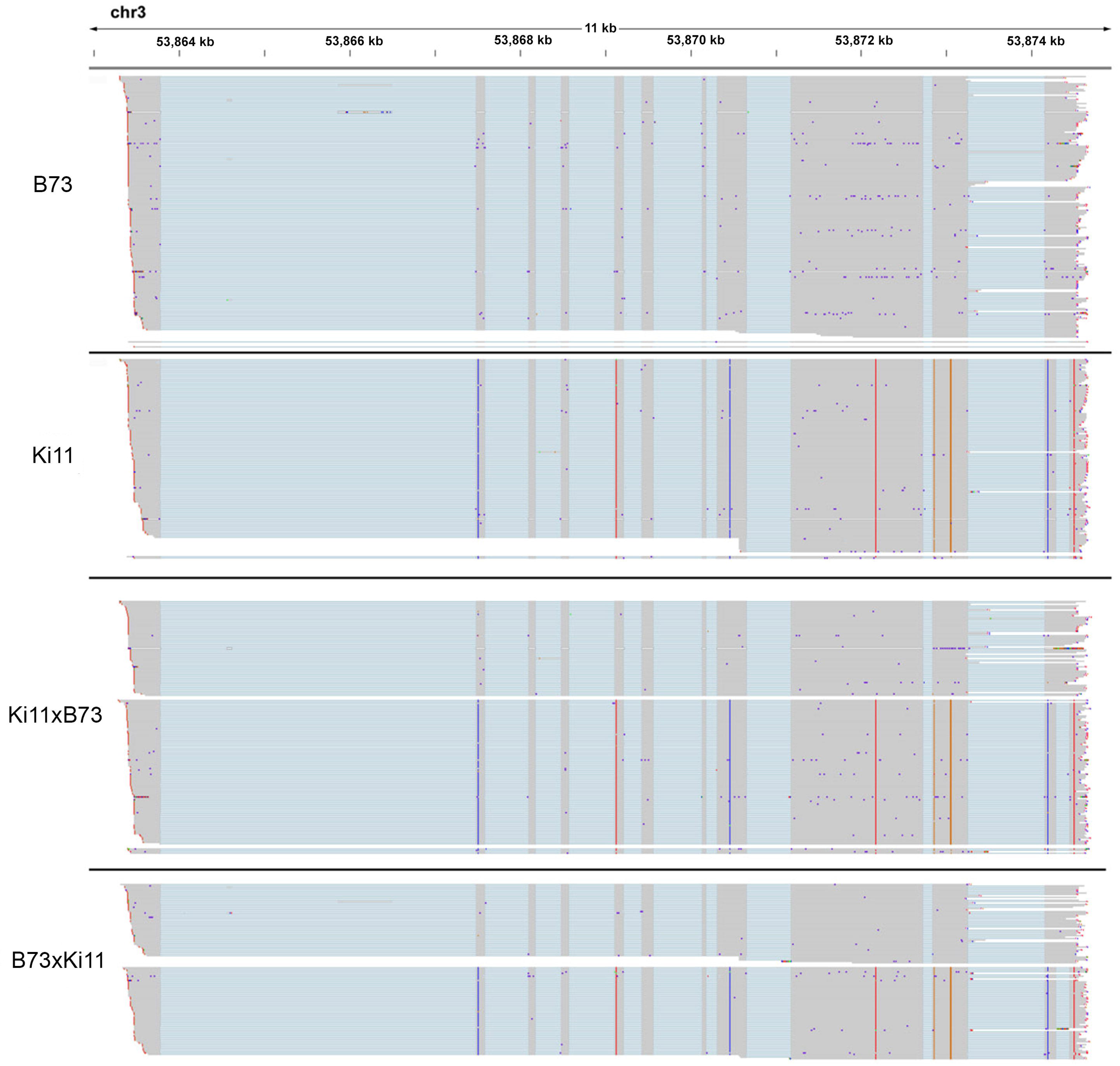
IsoPhase identifies allelic-specific isoform expression. The gene PB.8517 (Zm00001d040612) shows allelic-specific expression. B73 predominantly expresses the PB.8517.4 isoform, which has an unspliced 3’ exon. Ki11 predominantly expresses the PB.8517.1 isoform, in which the last 3’ exon spliced. The two F1 hybrids expresses both isoforms, but each isoform is associated with the parental allele.

We conclude that while short-read data achieves higher sequencing depth and can call more SNPs, full-length transcripts deliver accurate haplotype information with high specificity and can be used to study allele-specific expression. In future work, combining the deep coverage of short read data with full-length long-read data should dramatically improve both the sensitivity and specificity of transcript haplotyping.

### Functional Annotation of the SNPs

We performed functional annotation of SNPs called by IsoPhase using the maize reference genome annotation v4. Among all SNPs, 24% were synonymous variants. Of the non-synonymous variants, 22,093 had potential large effects on the function of 5,140 genes, including 21,685 missense, 287 splice donor/acceptor and 243 stop gained/retained variations. In addition, 10% and 17% variants were in 3’ UTRs and 5’ UTRs, respectively (Fig. 6a-b). Among those, 2,556 genes had SIFT (sort intolerant from tolerant) scores < 0.05 and were therefore predicted to be deleterious mutations. Gene ontology analysis revealed that most of the large-effect genes were associated with ‘molecular function’ terms related to catalytic activity and binding, and ‘biological process’ terms related to metabolic and cellular processes (Fig. 6c). The resultant differences in these processes could contribute to the phenotypic differences between Ki11 and B73, as well as the differences between the hybrids and their parents. We also found that these large-effect genes had more isoforms in root and embryo tissues in B73, but more isoforms in endosperm tissue in Ki11. In addition, the number of isoforms in the two reciprocal hybrids exhibited the high-parent value in root and embryo, but not in endosperm, suggesting that genes in endosperm play a role in determining the developmental differences between B73 and Ki11 plants (Fig. 6d-f)), the isoform-level agreement in embryo, endosperm, and root tissue had kappa scores of 0.935, 0.944, and 0.949, respectively, indicating good agreement between long-reads and short-reads.

**Figure 6:**
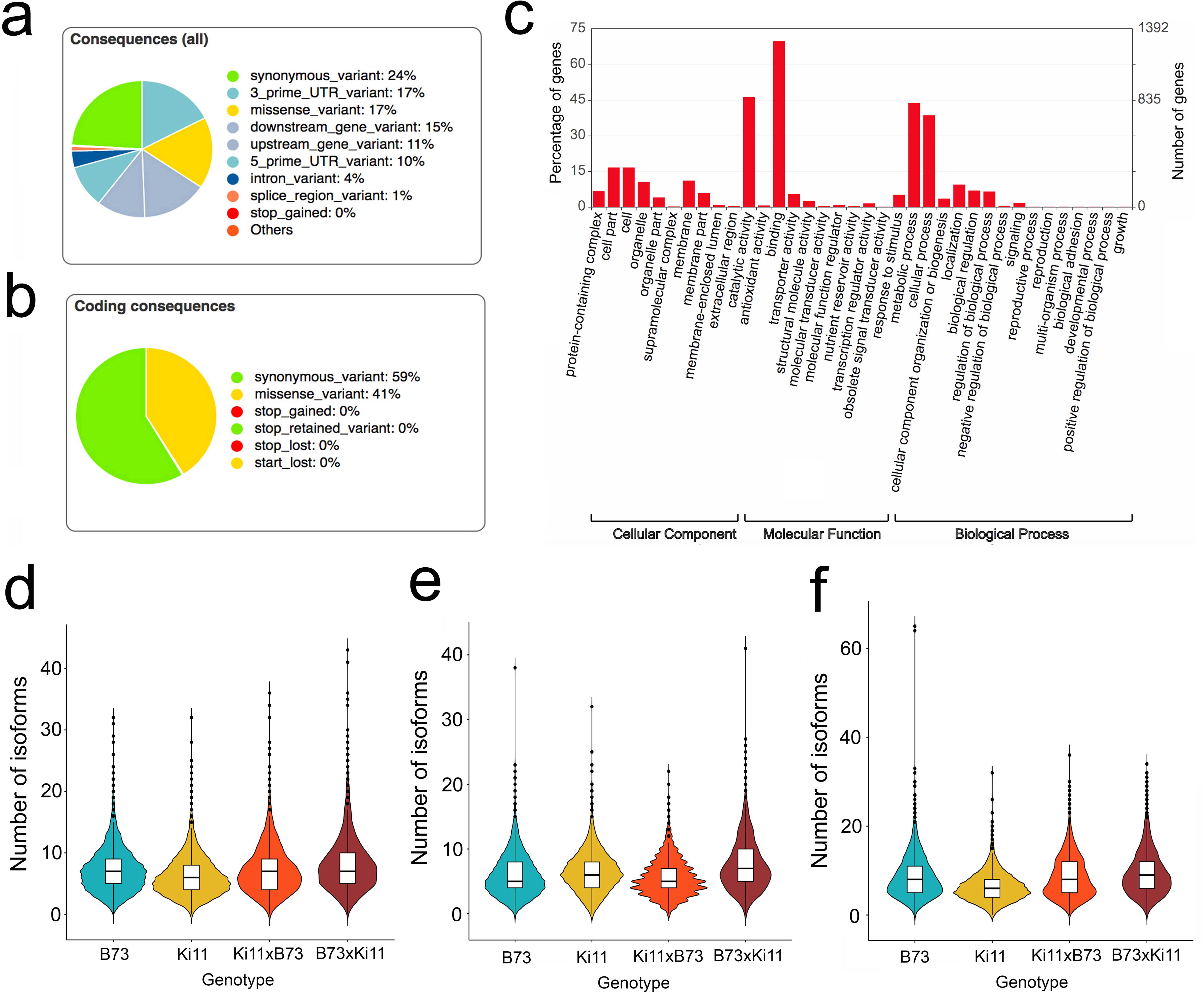
Annotations of the SNPs called from IsoPhase. a) Distribution of different categories of SNPs from variant effect predictor. b) Proportion of different categories of SNPs from coding sequences. c) Gene ontology analysis of large-effect genes. d-f) Number of isoforms of large-effect genes in two parents and reciprocal hybrids from root (d), endosperm (e) and embryo (f).

### Imprinted genes and cis-, trans-regulatory effects

We identified a number of imprinted genes based on the phasing results from IsoPhase, which were confirmed by short-read data (Supplementary Table 8, Supplementary Data 2). We found 26 paternally inherited genes in endosperm, of which 70% were the same as those identified in a previous study^18^; and 2 in embryo, which were also imprinted in endosperm. On the other hand, we found 30 maternally inherited genes in endosperm, of which 40% were the same as in the previous study, and 3 in embryo, of which two were also imprinted in endosperm. We found no imprinted genes in root. The expression levels of each allele confirmed the paternal and maternal expression pattern of these imprinted genes (Fig. 7a-b). Gene ontology analysis revealed that maternally expressed genes (MEGs) were slightly enriched in biological and molecular functions relate to establishment of localization, localization, and binding (Fig 7c). Paternally expressed genes (PEGs) were enriched in developmental process, multicellular organism process, reproduction, biological regulation, catalytic activity, and binding (Fig 7d), consistent with the previous study^18^.

**Figure 7:**
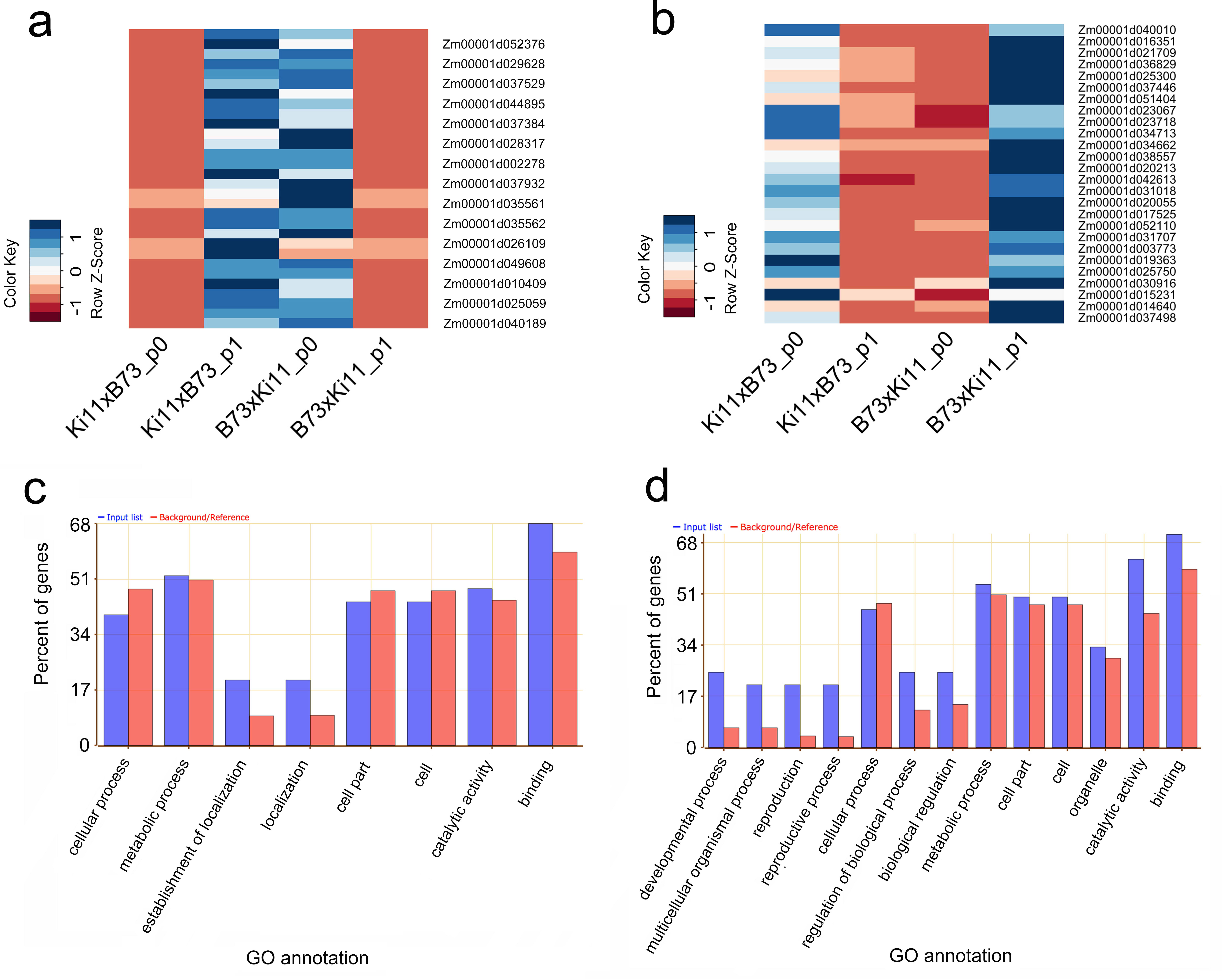
Allelic expression and gene ontology enrichment of maternal and paternal imprinted genes identified from phasing. (a-b) Allelic expression of maternal and paternal imprinted genes in reciprocal hybrids in endosperm. p0: phase0, which is B73 allele; p1: phase1, which is Ki11 allele. Data were normalized by Z score to compare expression change in different samples. (c–d) Gene ontology enrichment of maternal and paternal imprinted genes. Input list: imprinted genes; Background/reference: all annotated genes.

Variation in cis- and trans-regulation can be distinguished by ratios of allelic expression in parents relative to the F1 interspecific hybrid or allotetraploid. In order to see cis and trans effects on divergence in gene expression, we calculated the allelic ratios in parents and reciprocal hybrids from the phasing results, and used this information to identify genes with cis or trans effects. The results revealed that cis + trans effects were predominant in the two hybrids among all three tissues, followed by cis × trans effects, and that conserved genes were the least common (Supplementary Fig. 11a); these concepts are defined in Methods. In addition, the number of isoforms was slightly higher in conserved genes than in those with cis + trans or cis × trans effects (Supplementary Fig. 11b).

## Discussion

Maize is an important diploid genetic model for elucidating transcriptional networks. Recently, full-length transcript sequencing using long-read technology has enabled us to characterize alternative splicing events and improve maize genome annotation^14, 19^. However, the general Iso-Seq algorithm ignores SNP-level information, focusing instead on identifying alternative splicing differences. Heterosis has been extensively studied in plants using transcriptome sequencing approaches, revealing differentially expressed genes and biased expressions between parents and hybrid lines^20,21,22^. The results of these studies have provided clues about the molecular mechanisms underlying plant heterosis^23^. To date, however, no study has characterized allelic expression at single-molecule full-length transcript level. Maize, with its high diversity, provides a good model for studying heterosis.

On the other hand, transcriptome sequencing has a wide variety of applications, including evaluation of differences in gene expression between tissues, conditions, etc. By contrast, allele-specific expression analysis evaluates expression differences between two parental alleles in their hybrids. Both differential and allele-specific expression analyses have been employed to study heterosis using traditional RNA-seq^24, 25^, but the short read lengths of this technique make it impossible to construct the parental origins at the full-length transcript level. Full-length, single-molecule sequencing provides an unprecedented allele-specific view of the haploid transcriptome. In this study, haplotype phasing using long reads allowed us to accurately calculate allele-specific transcript and gene expression, as well as identify imprinted genes and investigate the cis/trans regulatory effects. Sequencing of full-length haplotype-specific isoforms enabled accurate assessment of allelic imbalance, which could be used to study the molecular mechanisms underlying genetic or epigenetic causative variants and associate expression polymorphisms with plant heterosis.

Maize is an excellent model for haplotype phasing studies: it is highly polymorphic; both inbreds and hybrids are easily obtainable; and a reference genomic sequence is available. In this study, we sequenced full-length cDNAs in two reciprocal hybrids and their parental lines using the PacBio Iso-Seq method. To phase the isoforms in hybrids, we developed IsoPhase, an accurate method for reconstructing haplotype-specific isoforms. Based on the assumption that the SNPs in each inbred line are homozygous, our method uses splice mapping to partition the reads into parental haplotypes. To our knowledge, in addition to IDP-ASE^6^ and IsoSeqASE (https://github.com/yunhaowang/IsoSeqASE) IsoPhase is another method capable of reconstructing the haplotype-specific isoforms from long single-molecule RNA-Seq data. Using IsoPhase, we successfully phased 6,907 genes using single-molecule sequencing data from maize reciprocal F1 hybrids and their parents. It is important to note that we only phased genes supported by more than 40 FL reads; the rest of the genes were not phased due to the sequencing depth cutoff. In addition, we used short-read data from the same tissues to confirm the SNPs called from long-read sequencing, giving us high confidence in the resultant phased genes. This is the first full-length isoform phasing study in maize, or in any plant, and thus provides important information for haplotype phasing to other organisms, including polyploid species. IsoPhase could also be used for self-incompatible species for haplotype phasing, although the high level of heterozygosity in such species would make this very challenging, and would also require deeper sequencing.

We observed that the allelic expression of phased genes varied among tissues. Among the phased genes, we also identified imprinted genes in endosperm and embryo, which were consistent with previous discovery. However, due to the large number of full-length reads required for high-confidence phasing by IsoPhase, future studies involving large-scale deep sequencing will be required to phase all genes in maize.

## METHODS

### Plant materials

Maize inbred lines B73 and Ki11 were grown at CSHL Uplands Farm, and reciprocal crosses were made between the two lines. For tissue collection, embryo and endosperm at 20 DAP (days after pollination) were collected for each genotype in two biological replicates, and root tissues were collected at 14 DAG (days after germination). All tissues were immediately frozen in liquid N_2_. For each tissue, at least 10 plants were pooled for each biological replicate.

### RNA preparation

Total RNA was prepared by grinding tissue in TRIzol reagent (Invitrogen 15596026) on dry ice and processing as recommended by the manufacturer. To remove DNA, an aliquot of total RNA was treated with RQ1 DNase (Promega M6101), followed by phenol/chloroform/isoamyl alcohol extraction and chloroform/isoamyl alcohol extraction using Phase Lock Gel Light tubes (5 PRIME 2302800), and ethanol precipitation. Precipitated RNA was stored at -20°C.

### Illumina RNA-Seq library construction

Total RNA (20 μg) was used for poly(A)^+^ selection using oligo(dT) magnetic beads (Invitrogen 610-02), eluted in water, and subjected to RNA-seq library construction using the ScriptSeq™ kit (Epicentre SS10906). Libraries were amplified by 15 cycles of PCR, and then sequenced in two lanes on the HiSeq 2500 PE150 platform at Woodbury Genome Center, Cold Spring Harbor Laboratory.

### PacBio library and single-molecule sequencing

cDNA was generated from 1 μg of total RNA per sample using the Clontech SMARTer PCR cDNA Synthesis Kit (catalog# 634925 or 634926) according to PacBio’s Iso-Seq Template Preparation for the Sequel System (https://www.pacb.com/wp-content/uploads/Procedure-Checklist-Iso-Seq-Template-Preparation-for-Sequel-Systems.pdf). Barcoded oligo-dT was conjugated to cDNAs during first-strand synthesis. The 16-bp barcode sequences used for this study are provided in Supplementary Table 1. The embryo, root, and endosperm cDNAs were enriched by PCR using PrimeSTAR GXL DNA Polymerase (Clontech, catalog# R050A or R050B). Amplification conditions used were as follows: initial denaturation at 98°C for 30 seconds, 9–12 cycles of 98°C for 10 seconds, 65°C for 15 seconds, and 68°C for 10 minutes. A final extension was performed at 68°C for 5 minutes. After amplification, tissues (embryo, root and endosperm) from the same strain were pooled in equimolar quantities, yielding four pools that were used to construct SMRTbell libraries using the SMRTbell Template Prep Kit 1.0 (Pacific Biosciences, Part No. 100-259-100). After library construction, each SMRTbell library was size-fractionated using SageELF (Sage Science). The final SMRTbell libraries were annealed with Sequencing Primer v4 (Pacific Biosciences Part No. 101-359-000) and bound with Sequel Binding Kit 2.1(Pacific Biosciences Part No. 101-429-300). The polymerase-bound SMRTbell libraries were loaded at 3–10 pM on-plate concentrations and sequenced using the Sequel Sequencing Kit 2.1 (Pacific Biosciences Part No. 101-312-100) and Instrument Software v5.1.

### Illumina data analysis

Raw reads were aligned to the B73 reference genome (RefGen_v4)^14^ using STAR 2.4.2a^26^ with minimum intron length of 20 bp and maximum intron length of 50 kb; default settings were used for the other parameters. Quantification of genes and isoforms was performed using cufflinks version 2.2.1^27^ using the GTF annotation file generated by PacBio sequencing.

### PacBio Data Analysis

PacBio data were analyzed by running the IsoSeq3 application in PacBio SMRT Analysis v6.0 to obtain high-quality, full-length transcript sequences, followed by downstream analysis.

### Full-Length Reads Classification

Full-length reads were identified as CCS reads that contained both the 5’ and 3’ primer and the poly(A) tail preceding the 3’ primer. The 5’ primer consists of the Clontech SMARTer cDNA primer with an ATGGG overhang. The 3’ primer consists of a tissue-specific 16-bp PacBio barcode followed by the Clontech SMARTer cDNA primer (Supplementary Table 1).

### Isoform-level Clustering Analysis to obtain High-Quality Transcript Sequences

To increase detection of rare isoforms, the demultiplexed FL reads were pooled to perform isoform-level clustering analysis^28^. After clustering, consensus sequences were called using the Arrow algorithm, and only polished sequences with predicted consensus accuracy ≥ 99% were considered high-quality (HQ) and retained for the next step.

### Mapping to B73 genome and filtering

The HQ transcript sequences were mapped to B73 v4 genome using minimap2 (version 2.11- r797)^14^ using parameters ‘-ax splice -t 30 -uf --secondary=no -C5’. We then filtered for alignments with ≥ 99% coverage and ≥ 95% identity and removed redundancy using scripts from cDNA_Cupcake http://github.com/Magdoll/cDNA_Cupcake). The full list of commands used at this step is as follows:

~~~
minimap2 -ax splice -uf -C5 --secondary=no B73_RefV4.fa hq.fastq > hq.fastq.sam
sort -k 3,3 -k 4,4n hq.fastq.sam > hq.fastq.sorted.sam
collapse_isoforms_by_sam.py --input hq.fastq --fq -s hq.fastq.sorted.sam \
     -c 0.99 -i 0.95 --dun-merge-5-shorter -o hq.no5merge
get_abundance_post_collapse.py hq.no5merge.collapsed cluster_report.csv
filter_away_subset.py hq.no5merge.collapsed
~~~

### Removing Potential Artifacts using SQANTI

We applied further filtering criteria to remove potential genomic contamination and rare PCR artifacts. We used a modified version of SQANTI^16^ that categorizes each isoform according to existing B73 RefGen_v4 annotations and a list of short-read junctions from the same samples. An isoform is retained in the dataset if: 1) it is FSM/ISM/NIC and does not have intra-priming; 2) it is NNC, does not have intra-priming, is not RT-switching, and all junctions are either all canonical or supported by short reads; or 3) it is antisense, intergenic, genic, does not have intra-priming, is not RT-switching, and all junctions are either all canonical or supported by short reads. The rationale behind the filtering is to eliminate artifacts that come from intra-priming (dT priming off genomic ‘A’ stretches), potential RT-switching, and other library or sequencing errors that could introduce erroneous splice junctions. Isoforms that are categorized as FSM/ISM/NIC are isoforms that use all known splice junctions, and are therefore trusted. For all other categories, we only retain the isoform if the junctions are either canonical or supported by short reads. This approach yields a high-confidence dataset that is well-supported by existing annotation and matching short-read data.

### De-multiplex Final Isoforms by Sample and Rarefaction Curves by Subsampling

We recovered the relative abundance of each the final isoforms in each sample by extracting the fraction of full-length reads supporting each isoform from each sample. To draw rarefaction curves, we used the unique Iso-Seq IDs (format: PB.X.Y) to indicate unique transcripts and the matching reference gene name (ex: Zm00001d027230) from B73 annotation. If a gene was novel, we created a novel gene ID (ex: novelGene_124) for each non-overlapping strand-specific locus. Subsampling was performed at 10,000 FL read intervals for 100 iterations, taking the average number of unique transcript/genes observed to plot the rarefaction curves. The observations of genotype specific genes/isoforms within each tissue were validated by short-read data from two biological replicates. Genes and isoforms were assigned to groups of genotypes according to whether they were expressed in each tissue. These assignments were compared to the long-read data assignments using Fleiss’ kappa^17^, a statistical measure that calculates the degree of agreement in classification over what would be expected by chance.

### Hybrid-inbred additive and nonadditive expression analysis

The additive and nonadditive expression of genes were identified based on previous method^3^. Briefly, the inbred expression values were used to calculate the expected additive expression level of each reciprocal hybrid. To account for the 2m:1p endosperm dosage, the expected additive values were calculated as: additive expression = [(2 × maternal expression level) + paternal expression level]/3. The expected additive values were compared with the appropriate reciprocal hybrid values for all two biological replications. A two-tailed homoscedastic t test was performed and all genes with P < 0.05 were considered to be nonadditively expressed. Both reciprocal hybrids were independently tested for each gene.

### SNP calling and Phasing using Iso-Seq data

All full-length reads from all 12 samples were aligned to the B73 RefGen_v4 genome using minimap2 (v2.11-r797)^15^ to create a pileup. Then, at each position with at least 10-base coverage, Fisher’s exact test with the Bonferroni correction was applied with a p-value cutoff of 0.01. Only substitution SNPs were called. Then, sample-specific haplotype information was obtained by looking at the number of FL reads associated with each allele. To account for residual errors in the FL reads, we error-corrected the haplotypes down to the two dominant haplotypes (maternal and parental haplotype) that minimizes the edit distance. Functional annotation of SNPs was performed using SnpEff^29^ and Gramene^30^ based on the maize B73 genome annotation RefGen_v4. SIFT 4G^31^ was used to predict deleterious mutations through the SciApps platform^32^. WEGO 2.0^33^ was used for the GO enrichment analysis of the large-effect genes under Pearson Chi-Square test.

### Discovering SNPs and measuring allelic expression

Short reads from all samples were mapped to the B73 genome using STAR 2.4.2a^26^. SAMtools^34^ was used to trim out reads for which mapping quality was less than 20. The alignments of each sample were merged together, pileuped by SAMtools, and used for calling SNPs. SNPs were called if a site was covered by at least 10 reads in both B73×Ki11 and Ki11×B73. The expression level of B73 alleles of each SNP site was determined by summary of the B73 reads that mapped to the SNP site, and the expression level of Ki11 alleles of each SNP site were calculated using the sum of the Ki11-reads of the SNP site.

### Identification of paternal and maternal imprinted genes

To identify the imprinted SNPs, we used the method of Zhang *et al*^18^. In brief, Bm represents the percentage of expression for maternal alleles out of all expression including the maternal and paternal alleles, whereas Bp represents the expressional percentage for the paternal alleles in B73 × Ki11 and Ki11 × B73 reciprocal hybrids. To obtain high-quality imprinted genes, we employed a very high stringent standard, such that a SNP was considered to be imprinted if the expression level of the actively expressed allele was at least five-fold higher than that of the imprinted repressed allele. Given that the endosperm has two doses of maternal alleles and only one dose of paternal alleles, therefore the maternally preferentially expressed SNPs should have Bm values of greater than 10/11 [Bm > 2×5 / (2×5 +1)], whereas the paternal preferentially expressed SNPs should have Bp value of greater than 5/7 {Bp > 5/ (2×1 +5)}. In the meantime, we applied this approach to the phased genes in this study, from which the full-length counts of each haplotype were used to calculate Bm and Bp values. As a final set of imprinted genes, we only chose those common to both datasets, which are presumed to be very high-confidence. agriGO v2.0^35^ was used for the GO enrichment analysis of the imprinted genes under Pearson Chi-Square test.

### Identification of cis- and trans-regulatory effects

Cis and trans effects were determined according to Bell *et al.,* 2013^36^. In brief, we compared the ratio of expression of the two parental alleles in F1 hybrids with the relative expression of the same alleles in the homozygous parents. Genes were categorized as “cis only” or “trans only” if parental expression divergence was observed as well as significant cis-effects or trans-effects, but not both. “Cis + trans” refers to cases where cis- and trans-effects were both significant and promoted expression of the same allele, whereas “cis × trans” indicates counteracting effects of cis- and trans-regulators. A “conserved” pattern of regulation was assigned to cases where there was no significant cis-effect, trans-effect, or parental expression divergence.

### Contributions

B.W., E.T., and D.W. conceived the idea for the study; B.W. collected the tissues; M.R. generated RNA and Illumina libraries; P.B. and K.E. generated the PacBio libraries and data; B.W., E.T., Y.J., L.W., A.O., and K.C. analyzed the data; and B.W. and E.T. wrote the manuscript.

### Accession codes

The data generated in this study, including PacBio Iso-Seq reads and Illumina short reads, have been submitted to ArrayExpress (https://www.ebi.ac.uk/arrayexpress/) under accession numbers E-MTAB-7837 and E-MTAB-7394. The IsoPhase tool developed in this study is available in the GitHub repository: https://github.com/magdoll/cdna_cupcake.

## Acknowledgements

This work was supported by USDA-SCA grant #51530311 and NSF Gramene grant #52930511. The authors thank Tim Mulligan for assistance with farming.

