## Supplemental information for "Variant Phasing and Haplotypic Expression from Single-molecule Long-read Sequencing in Maize"

**Supplementary Table 1. 16-mer Barcodes corresponding to the Sample.**

| Sample | Genotype | Barcode Sequence |
|---|---|---|
| Embryo 1 | B73 20 DAP | TCAGACGATGCGTCAT |
| Embryo 2 | Ki11 20 DAP | TACTAGAGTAGCACTC |
| Embryo 3 | Ki11xB73 20 DAP | GATCTCTACTATATGC |
| Embryo 4 | B73xKi11 20 DAP | CATAGCGACTATCGTG |
| Endosperm 1 | B73 20 DAP | CTATACATGACTCTGC |
| Endosperm 2 | Ki11 20 DAP | TGTGTATCAGTACATG |
| Endosperm 3 | Ki11xB73 20 DAP | ACAGTCTATACTGCTG |
| Endosperm 4 | B73xKi11 20 DAP | CGAGCACGCGCGTGTG |
| Root 1 | B73 20 DAP | GCTCGACTGTGAGAGA |
| Root 2 | Ki11 20 DAP | TGCTCGCAGTATCACA |
| Root 3 | Ki11xB73 20 DAP | TCACACTCTAGAGCGA |
| Root 4 | B73xKi11 20 DAP | CGCTGCGAGAGACAGT |

**Supplementary Table 2. Number of full-length, non-concatemer (FLNC) reads from each of the 12 samples after demultiplexing.**

| SAMPLE | NAME | # of FLNC |
|---|---|---|
| EM1 | B73 Embryo | 339,048 |
| EM2 | Ki11 Embryo | 254,342 |
| EM3 | Ki11xB73 Embryo | 305,307 |
| EM4 | B73xKi11 Embryo | 444,580 |
| END1 | B73 Endosperm | 284,678 |
| END2 | Ki11 Endosperm | 290,122 |
| END3 | Ki11xB73 Endosperm | 232,168 |
| END4 | B73xKi11 Endosperm | 288,205 |
| R1 | B73 Root | 362,431 |
| R2 | Ki11 Root | 225,208 |
| R3 | Ki11xB73 Root | 287,485 |
| R4 | B73xKi11 Root | 426,238 |
| **TOTAL** | | **3,739,812** |

**Supplementary Table 3. Mapping high-quality (HQ) transcript sequences to the genome and filtering criteria.** HQ sequences were mapped to B73 v4 genome and filtered for 99% coverage and 95% identity. Redundant transcripts are collapsed.

| **Mapping to Genome** | |
|---|---|
| HQ Isoforms | 250,168 |
| Mapped to maize v4 | 248,424 (99.3%) |
| Mapped to maize v4 with 99% coverage 95% identity | 229,757 (91.8%) |
| **After collapse & compare to annotation:** | |
| Number of unique isoforms | 90,419<br>(27,967 loci) |

**Supplementary Table 4. Removal of library artifacts using a modified version of the SQANTI software.**

| **Cause of SQANTI Filtering** | **Transcripts Filtered** |
|---|---|
| **BEFORE FILTERING:** | 90,419 transcripts<br>(27,967 loci) |
| Intrapriming | 1402 |
| RT-switching | 1906 |
| Some junctions not canonical and not all junctions short-read-supported | 11993 |
| **AFTER FILTERING:** | 75,118 transcripts<br>(23,412 loci) |

**Supplementary Table 5. Top BLASTN hit counts of the unmapped high-quality (HQ) transcript sequences to the NR database.** BLASTN was run with a report of best hit with E-value cutoff of 0.1; 1669 of 1744 HQ sequences had a BLASTN hit.

| Species | Sequence_Hit_Counts |
|---|---|
| Bipolaris zeicola | 534 |
| Zea mays | 523 |
| Cochliobolus sativus | 200 |
| Fusarium verticillioides | 162 |
| Bipolaris maydis | 63 |
| Trichoderma virens | 39 |
| Bipolaris victoriae | 17 |
| Setosphaeria turcica | 9 |

**Supplementary Table 6. Sample-specific transcript counts.** Using the demultiplexed full-length reads, we assigned Iso-Seq transcripts back to each sample. If a transcript contained at least one full-length read from a sample, it was considered to be expressed.

| Sample | Number |
|---|---|
| B73-Embryo | 31,767 |
| B73-Endosperm | 25,620 |
| B73-Root | 32,649 |
| Ki11-Embryo | 25,024 |
| Ki11-Endosperm | 24,773 |
| Ki11-Root | 24,281 |
| Ki11xB73-Embryo | 29,287 |
| Ki11xB73-Endosperm | 23,704 |
| Ki11xB73-Root | 30,331 |
| B73xKi11-Embryo | 33,854 |
| B73xKi11-Endosperm | 25,677 |
| B73xKi11-Root | 34,820 |

#### Supplementary Table 7. Fleiss' kappa of gene-level agreement and isoform-level agreement in embryo, endosperm, and root tissue between long and short reads.

| Tissue | Gene | Isoform |
|---|---|---|
| embryo | 0.905 | 0.958 |
| endosperm | 0.915 | 0.964 |
| root | 0.926 | 0.966 |

Genes and isoforms were assigned to groups of genotypes according to whether they were expressed in each tissue. These assignments were compared to the long-read data assignments using Fleiss' kappa, a statistical measure that calculates the degree of agreement in classification over what would be expected by chance.

#### Table S8. List of imprinted genes in 20DAP endosperm and embryo.

| RefGen_V4 Gene ID | Annotation | Imprint Type | Confirmed imprinting | RefGen_V3 Gene ID | Tissue |
|---|---|---|---|---|---|
| Zm00001d002278 | OSJNBa0058K23.15 protein; protein | MEG | NO | GRMZM2G107711 | Endosperm |
| Zm00001d004401 | Germin-like protein subfamily 1 member 8 | MEG | YES | GRMZM2G045809 | Endosperm |
| Zm00001d004707 | thylakoid assembly1 | MEG | NO | GRMZM2G090086 | Endosperm |
| Zm00001d009908 | UDP-glucuronic acid decarboxylase 4 | MEG | NO | GRMZM2G007404 | Endosperm |
| Zm00001d010409 | Peroxidase 29 | MEG | YES | GRMZM2G150134 | Endosperm |
| Zm00001d010451 | GDSL esterase/lipase | MEG | NO | GRMZM2G044947 | Endosperm |
| Zm00001d016394 | OSJNBa0087H01.6 protein; protein | MEG | NO | GRMZM2G021998 | Endosperm |
| Zm00001d023680 | Pollen Ole e 1 allergen and extensin family protein | MEG | NO | GRMZM2G048175 | Endosperm |
| Zm00001d024876 | Coatomer subunit beta-1 | MEG | YES | GRMZM2G116626 | Endosperm |
| Zm00001d025059 | Germin-like protein subfamily 1 member 8 | MEG | NO | GRMZM2G166141 | Endosperm |
| Zm00001d026109 | Putative polyphenol oxidase family protein | MEG | NO | AC209206.3_FG014 | Endosperm |
| Zm00001d027593 | proteasome component3 | MEG | NO | GRMZM2G472167 | Endosperm |
| Zm00001d028317 | Leucine-rich repeat receptor-like serine/threonine-protein kinase BAM3 | MEG | NO | GRMZM2G043584 | Endosperm |

| | | | | | |
|---|---|---|---|---|---|
| Zm00001d028398 | Glycosyl hydrolase family 10 protein | MEG | NO | GRMZM2G108032 | Endosperm |
| Zm00001d029628 | cystatin7 | MEG | NO | GRMZM2G148925 | Endosperm |
| Zm00001d030314 | Proline-rich protein | MEG | YES | GRMZM2G114356 | Endosperm |
| Zm00001d033528 | Protein NRT1/ PTR FAMILY 2.9 | MEG | YES | GRMZM2G104542 | Endosperm |
| Zm00001d033585 | leaf permease1 | MEG | YES | GRMZM5G858417 | Endosperm |
| Zm00001d035561 | Expressed protein; Mannose-specific jacalin-related lectin; protein | MEG | YES | GRMZM2G050412 | Endosperm |
| Zm00001d035562 | Expressed protein; Mannose-specific jacalin-related lectin; protein | MEG | NO | GRMZM2G163406 | Endosperm |
| Zm00001d037209 | Probable RNA-binding protein ARP1 | MEG | YES | GRMZM2G073700 | Endosperm |
| Zm00001d037384 | Anthocyanidin 3-O-glucosyltransferase | MEG | YES | GRMZM2G383404 | Endosperm |
| Zm00001d037529 | Acyl-CoA N-acyltransferase with RING/FYVE/PHD-type zinc finger protein | MEG | NO | GRMZM2G402538 | Endosperm |
| Zm00001d037932 | Transducin/WD40 repeat-like superfamily protein | MEG | NO | GRMZM2G325804 | Endosperm |
| Zm00001d040189 | no-apical-meristem-related protein1 | MEG | YES | GRMZM2G062650 | Endosperm |
| Zm00001d044340 | aldehyde dehydrogenase5 | MEG | NO | GRMZM2G097706 | Endosperm |
| Zm00001d044895 | ATPase%2C coupled to transmembrane movement of substance | MEG | NO | GRMZM5G874756 | Endosperm |
| Zm00001d049608 | Polycomb group protein FIE1 | MEG | YES | GRMZM2G118205 | Endosperm |
| Zm00001d051923 | GDSL esterase/lipase LTL1 | MEG | YES | GRMZM2G384780 | Endosperm |
| Zm00001d052376 | protein kinase homolog2 | MEG | NO | GRMZM2G099754 | Endosperm |
| Zm00001d003773 | OTU-like cysteine protease family protein | PEG | NO | GRMZM2G045185 | Endosperm |
| Zm00001d014640 | E3 ubiquitin-protein ligase BRE1-like 1 | PEG | NO | GRMZM2G162184 | Endosperm |
| Zm00001d015231 | Glucan endo-1%2C3-beta-glucosidase 6 | PEG | YES | GRMZM2G097207 | Endosperm |
| Zm00001d016351 | Eukaryotic initiation factor 4A-2 | PEG | YES | GRMZM2G028366 | Endosperm |
| Zm00001d017525 | Mitogen-activated protein kinase kinase kinase YODA | PEG | YES | AC209208.3_FG001 | Endosperm |
| Zm00001d019363 | Serine/threonine protein phosphatase 2A 59 kDa regulatory subunit B' eta isoform | PEG | YES | GRMZM2G106141 | Endosperm |
| Zm00001d020055 | Polyadenylate-binding protein-interacting protein 9 | PEG | NO | GRMZM2G093947 | Endosperm |
| Zm00001d020213 | Probable receptor-like protein kinase | PEG | YES | GRMZM2G132184 | Endosperm |

| | | | | | |
|---|---|---|---|---|---|
| Zm00001d021709 | isopentenyl pyrophosphate isomerase1 | PEG | YES | GRMZM2G108285 | Endosperm |
| Zm00001d023067 | unknown | PEG | NO | NO_V3_ID | Endosperm |
| Zm00001d023718 | defective18 | PEG | YES | GRMZM2G091819 | Endosperm |
| Zm00001d025300 | Sucrose nonfermenting 4-like protein | PEG | YES | GRMZM5G845175 | Endosperm |
| Zm00001d025750 | Probable E3 ubiquitin-protein ligase ARI8 | PEG | YES | GRMZM2G006428 | Endosperm |
| Zm00001d030916 | Extra-large guanine nucleotide-binding protein 1 | PEG | NO | GRMZM2G127739 | Endosperm |
| Zm00001d031018 | Ubiquitin-like-specific protease 1D | PEG | NO | GRMZM2G072939 | Endosperm |
| Zm00001d031707 | La-related protein 6B | PEG | YES | GRMZM2G045503 | Endosperm |
| Zm00001d034662 | Protein kinase APK1A chloroplastic | PEG | YES | GRMZM2G068117 | Endosperm |
| Zm00001d034713 | Anthranilate synthase alpha subunit 2 chloroplastic | PEG | YES | GRMZM2G138382 | Endosperm |
| Zm00001d036829 | alpha/beta-Hydrolases superfamily protein | PEG | YES | GRMZM2G092895 | Endosperm |
| Zm00001d037446 | Protein decapping 5 | PEG | YES | GRMZM2G341027 | Endosperm |
| Zm00001d037498 | L-tryptophan--pyruvate aminotransferase 1 | PEG | YES | GRMZM2G127160 | Endosperm |
| Zm00001d038557 | nuclear factor Y subunit C11 | PEG | YES | GRMZM2G440949 | Endosperm |
| Zm00001d040010 | unknown | PEG | YES | GRMZM2G121683 | Endosperm |
| Zm00001d042613 | RING/FYVE/PHD-type zinc finger family protein | PEG | YES | GRMZM2G025703 | Endosperm |
| Zm00001d051404 | Putative MAPKKK family protein kinase isoform 1%3B Putative MAPKKK family protein kinase isoform 2 | PEG | YES | GRMZM2G093316 | Endosperm |
| Zm00001d052110 | opaque endosperm1 | PEG | NO | GRMZM2G449909 | Endosperm |
| Zm00001d010409 | Peroxidase 29 | MEG | YES | GRMZM2G150134 | Embryo |
| Zm00001d025059 | Germin-like protein subfamily 1 member 8 | MEG | NO | GRMZM2G166141 | Embryo |
| Zm00001d040189 | no-apical-meristem-related protein1 | MEG | YES | GRMZM2G062650 | Embryo |
| Zm00001d016351 | Eukaryotic initiation factor 4A-2 | PEG | YES | GRMZM2G028366 | Embryo |
| Zm00001d037498 | L-tryptophan--pyruvate aminotransferase 1 | PEG | YES | GRMZM2G127160 | Embryo |

### Supplementary Figures

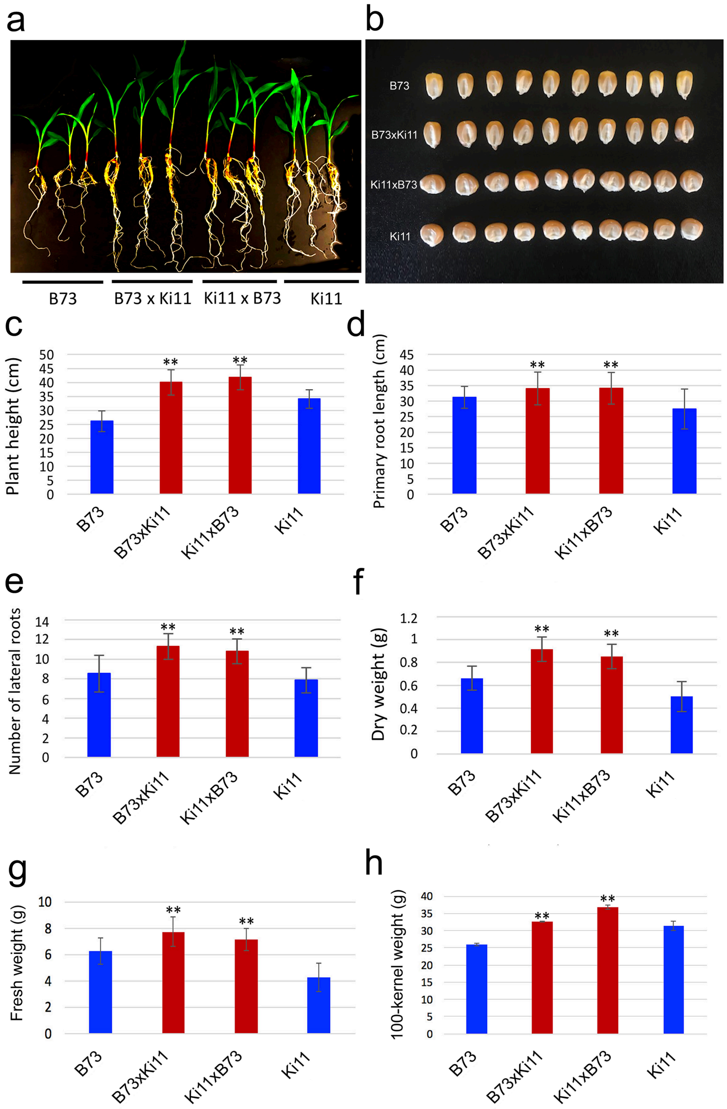


**Supplementary Fig. 1. Phenotype of maize B73, Ki11, and two reciprocal hybrids (B73 × Ki11, Ki11 × B73) at 14DAG and measure of different traits between parents and the hybrids.** (a) Shoot and root phenotype of B73, Ki11, and the two reciprocal hybrids. (b) Seed phenotype of B73, Ki11, and the two hybrids. (c) Plant height, (d) primary root length, (e) lateral root number, (f–g) biomass, and (h) 100-kernel weight of the parents and hybrids. ** $p<0.01$.

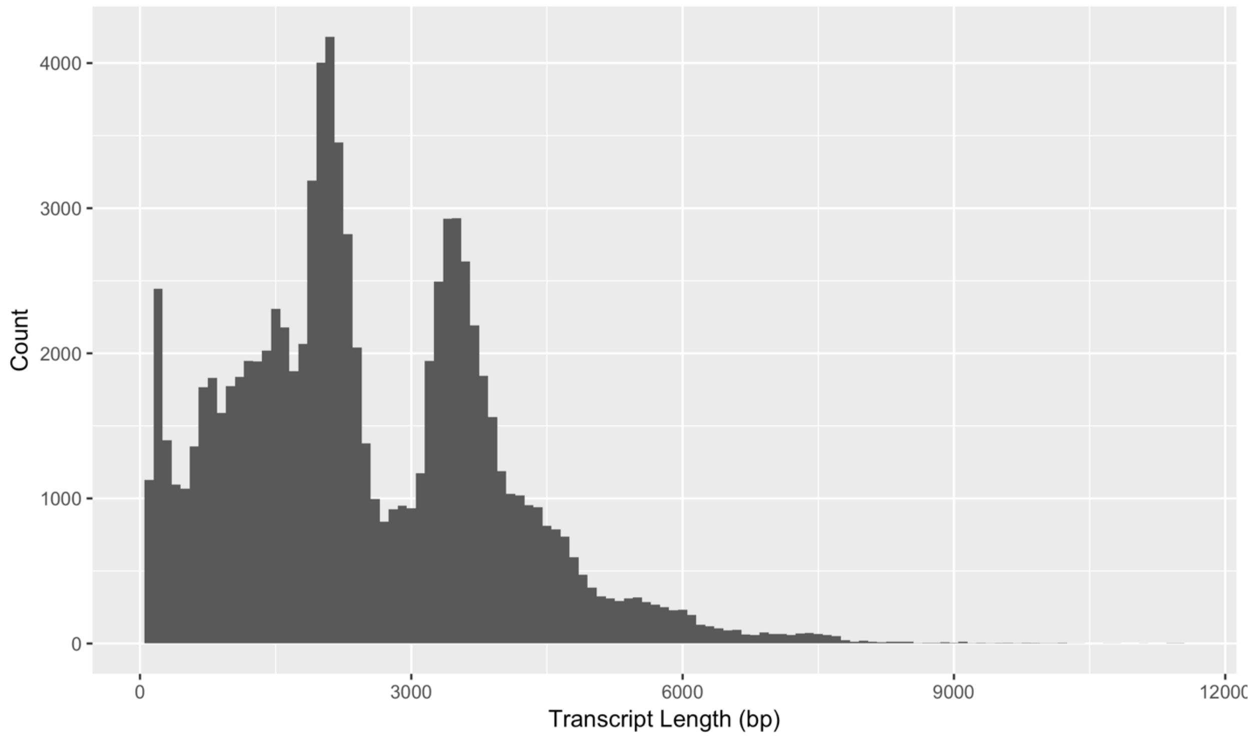


**Supplementary Fig. 2 Length distribution of the final transcript set.**

After mapping the high-quality (HQ) sequences to the B73 RefGen_v4 genome and filtering for coverage, identity, and running the SQANTI software to remove library artifacts, we obtain 75,118 transcripts. Min: 80bp, Max: 11,495 bp, Mean: 2,482 bp, $5^{th}$-$95^{th}$ percentile: 363-4,975 bp.

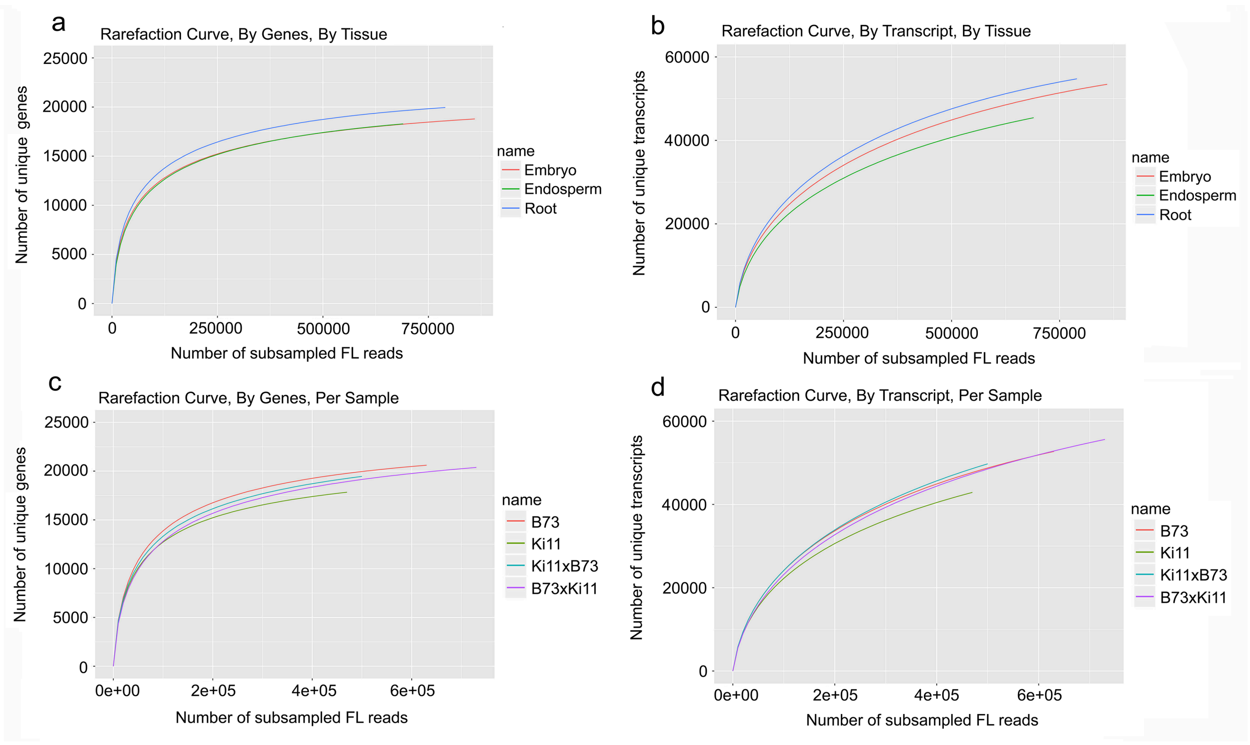


**Supplementary Fig. 3 Rarefaction curves against known genes and transcripts (a-b) by strain, (c-d) by tissue.** For each subpanel, the X-axis shows the number of subsampled full-length reads and the Y-axis shows the number of observed unique genes or transcripts. Both known and novel genes/transcripts are considered.

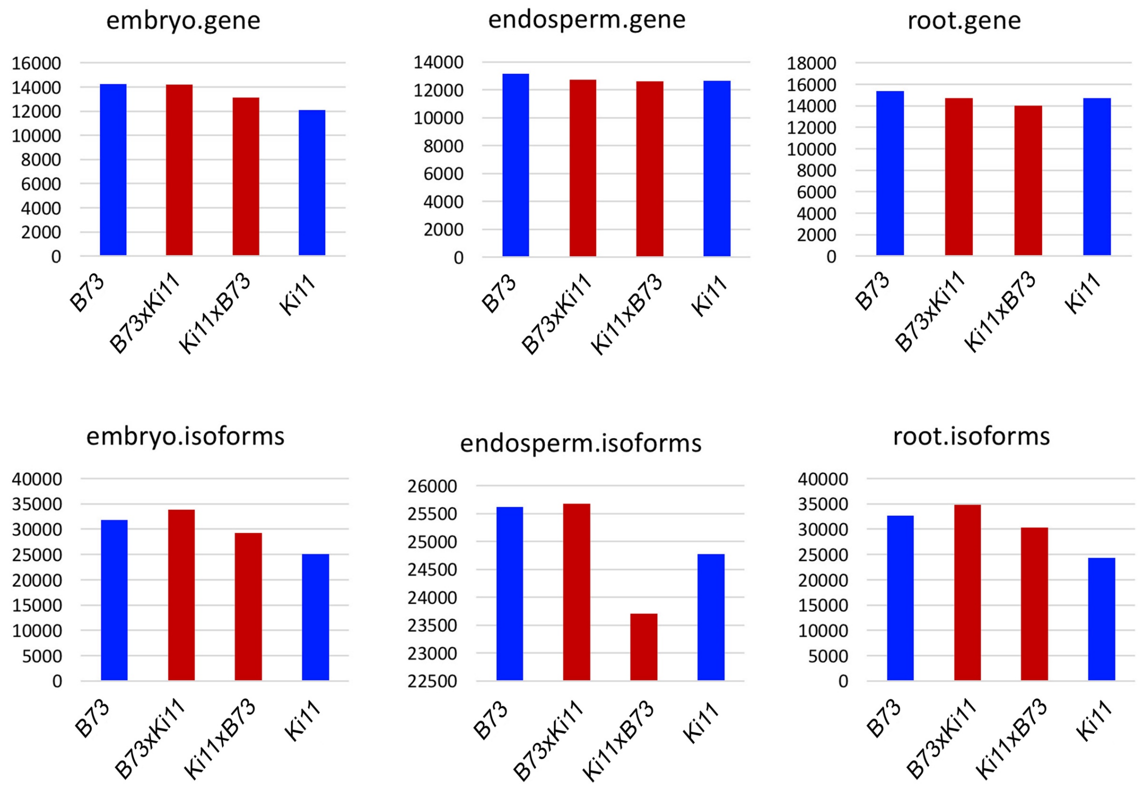


**Supplementary Fig. 4 Number of genes and isoforms between parents and two reciprocal hybrids in embryo, endosperm and root tissues.**

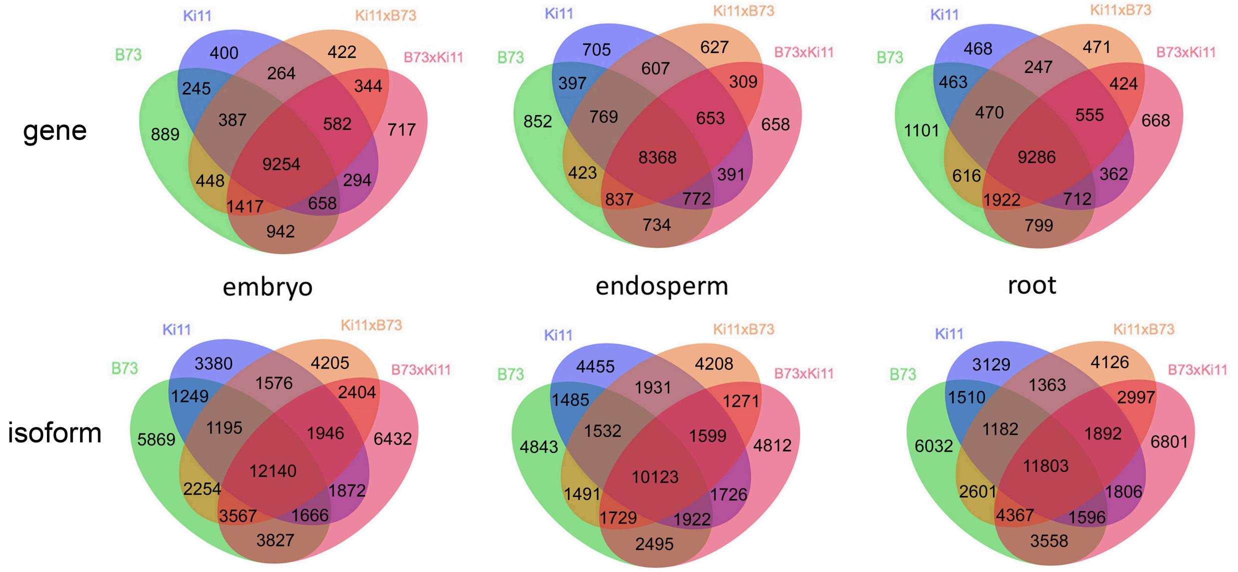


**Supplementary Fig. 5 Overlap of genes and isoforms among parents and two hybrids in embryo, endosperm and root.**

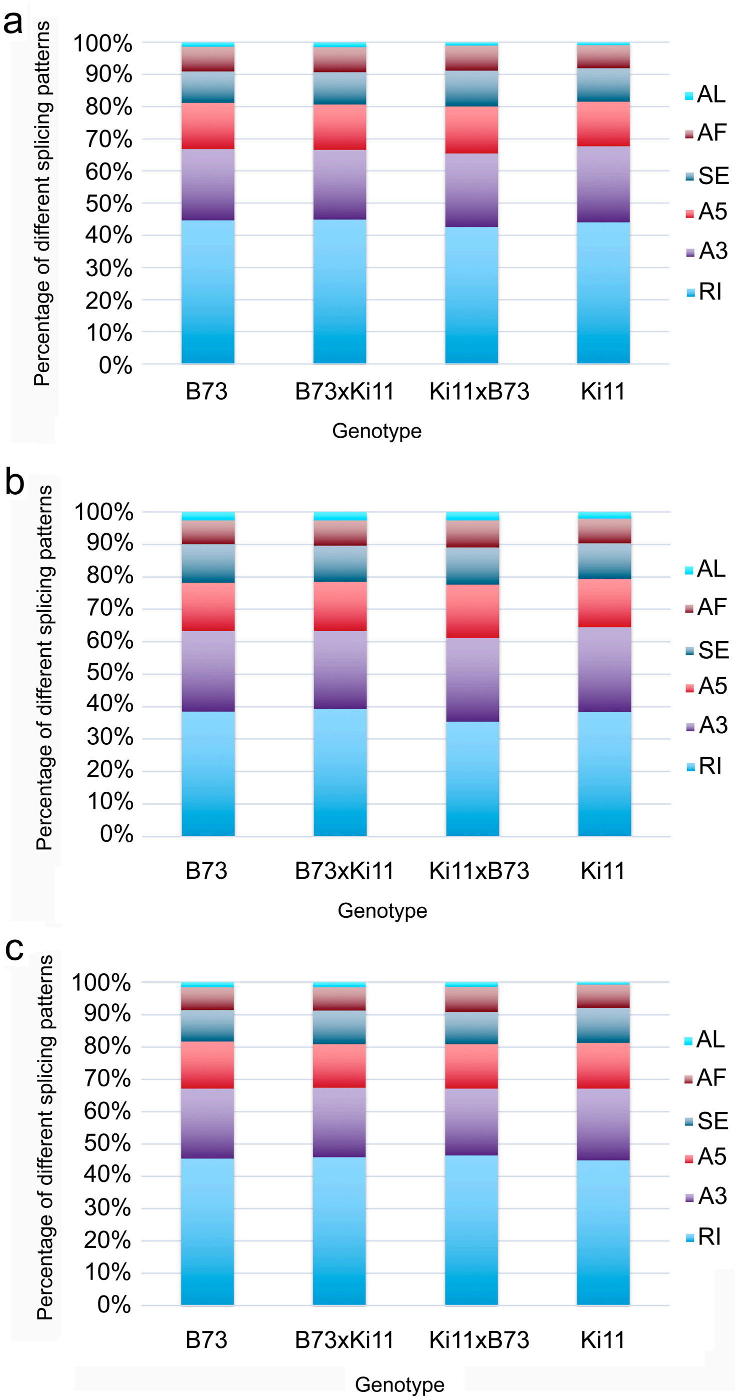


**Supplementary Fig. 6 Alternative splicing pattern between parents and two reciprocal hybrids in embryo (a), endosperm (b) and root (c).** A3: Alternative 3’ splice site; A5: Alternative 5’ splice site; AF: Alternative first exon’ AL: Alternative last exon; RI: Retained Intron; SE: Skipped exon.

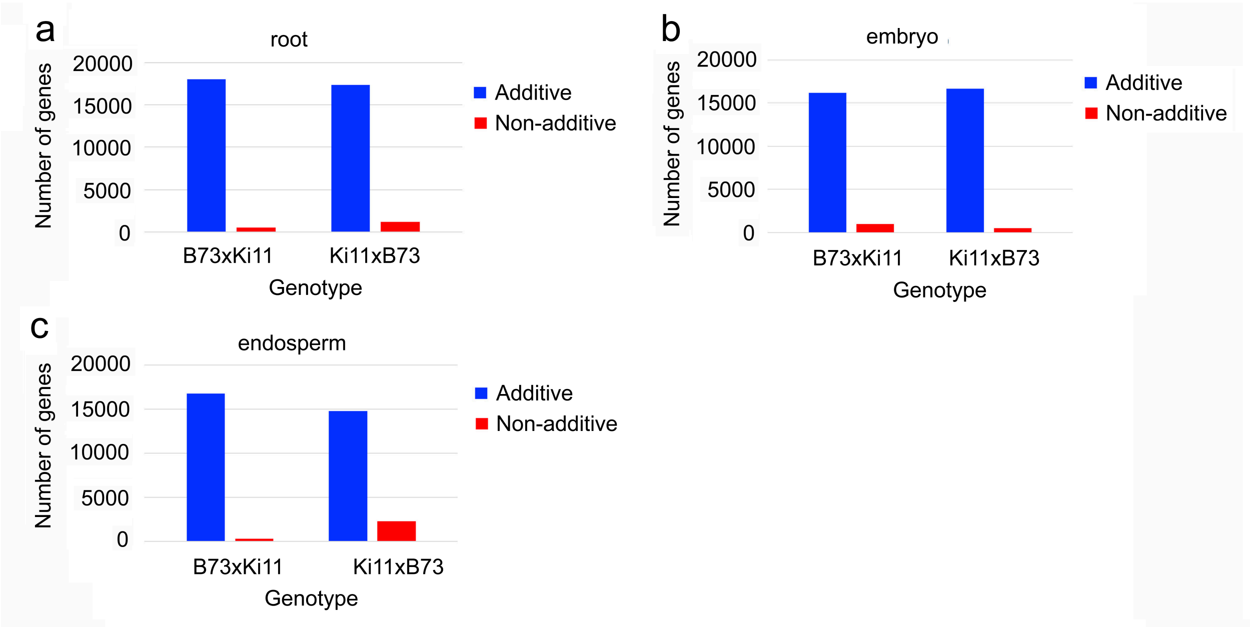


**Supplementary Fig. 7 Number of additive and non-additive expression genes in root (a), embryo (b) and endosperm (c).**

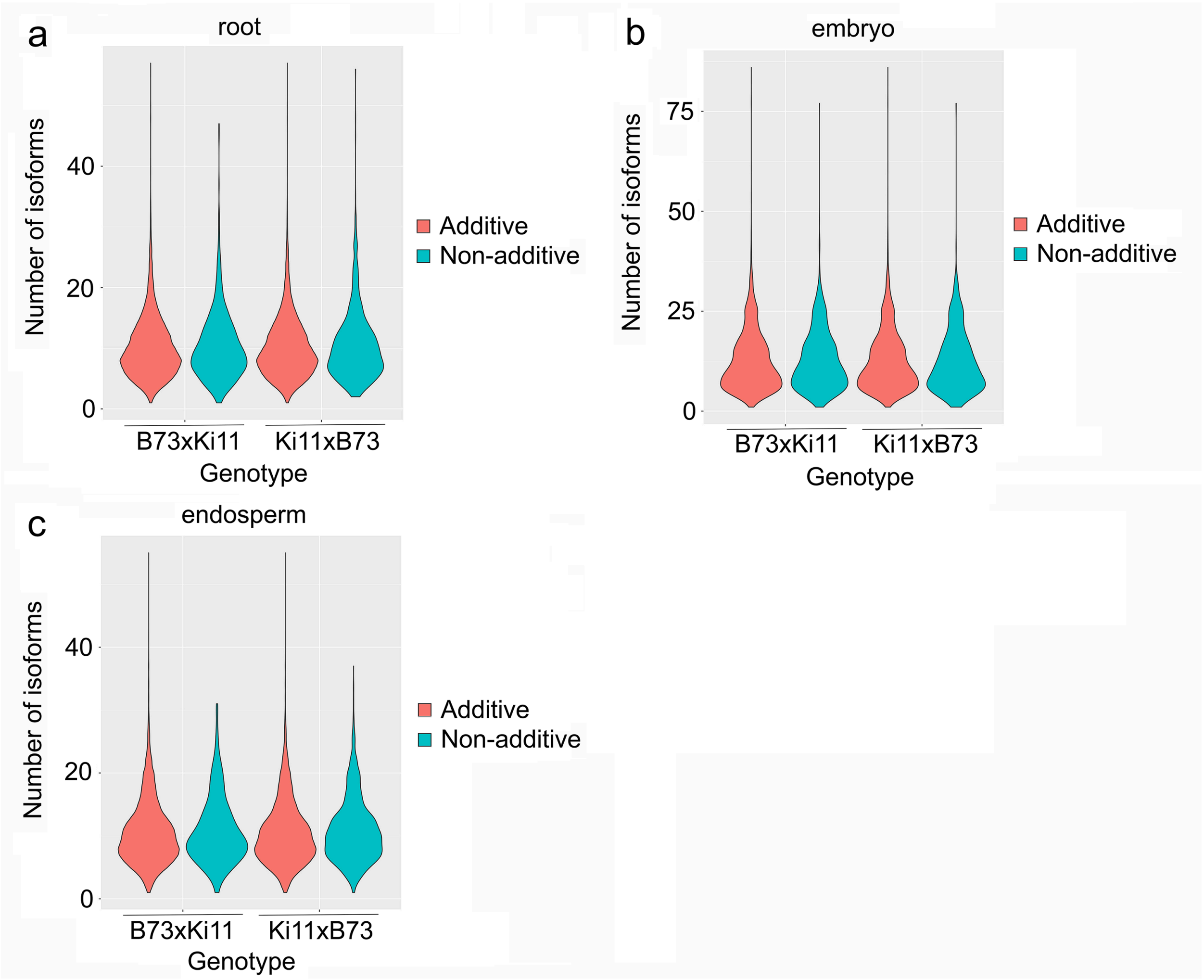


**Supplementary Fig. 8 Number of isoforms of additive and non-additive expression genes in root (a), embryo (b) and endosperm (c).**

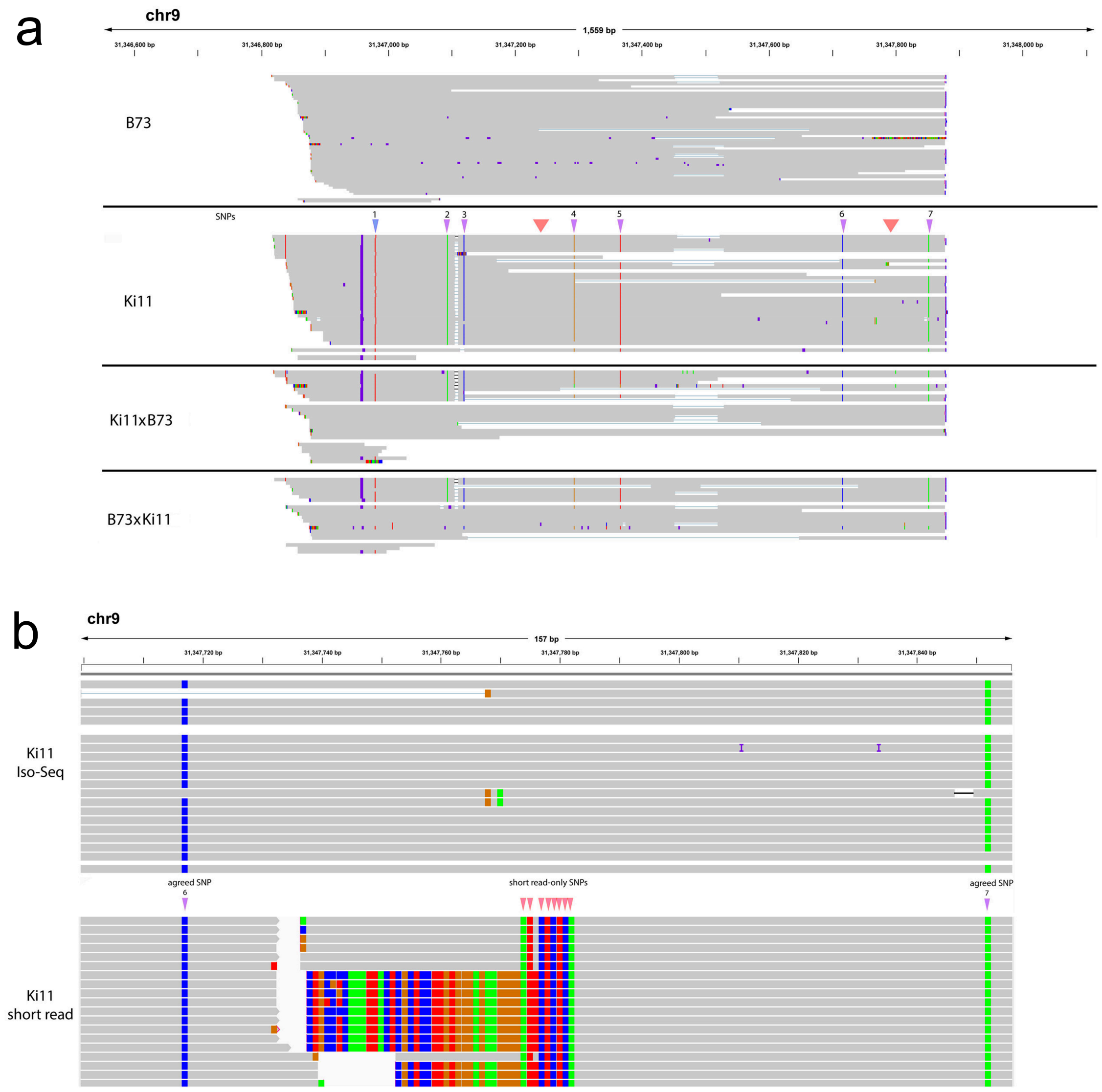


**Supplementary Fig. 9 IsoPhase phasing example.** a) The gene PB.21897 (Zm00001d045657) phased by IsoPhase. SNPs are depicted between the B73 and Ki11 tracks, seven SNPs are shown. SNP #2–#7 were called by based on both long- and short-read data (purple), SNP #1 was missed by long-read data due to reduced coverage (blue); suspicious short read-only SNPs are marked in red. b) Zoomed-in region between SNP #6 and SNP #7 showing suspicious SNPs (red) called based on short-read only. Top track shows Ki11 Iso-Seq FL reads; bottom track shows Ki11 short reads.

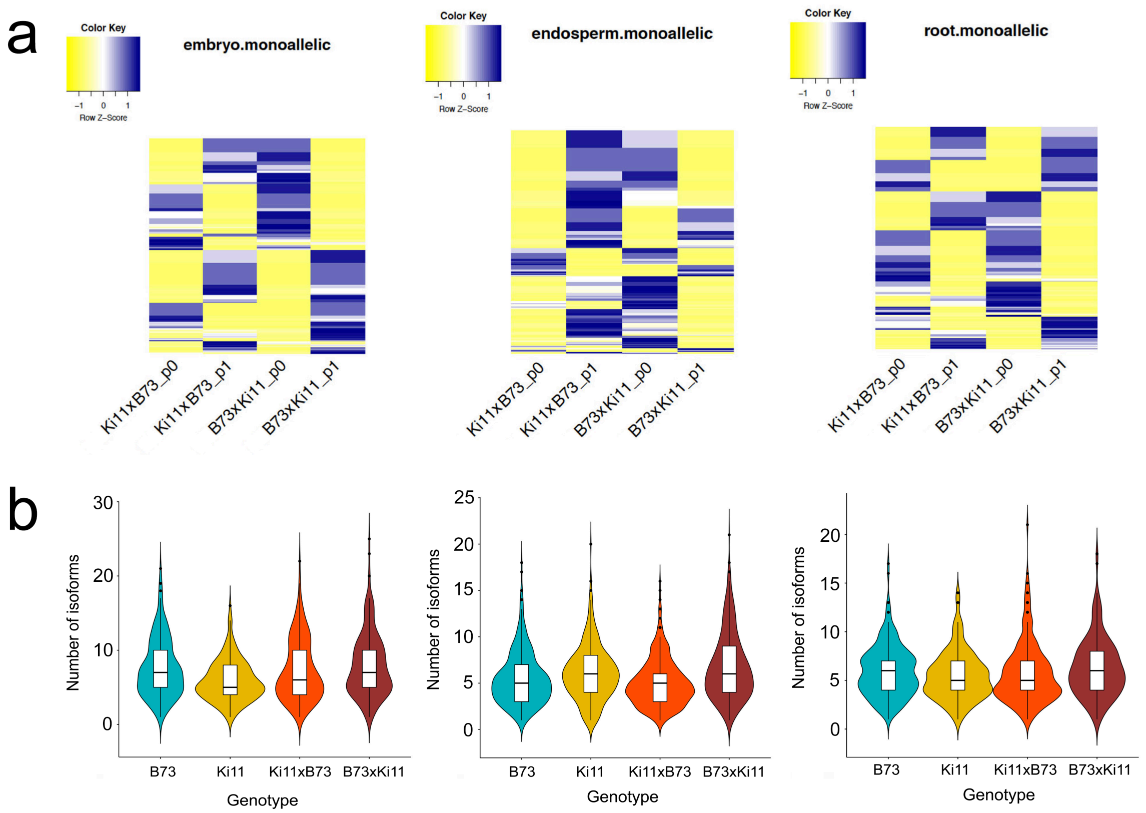


**Supplementary Fig. 10 Expression and number of isoforms of mono-allelic genes in different tissues.** a) Allelic expression of mono-allelic genes in reciprocal hybrids, and b) number of isoforms between parents and hybrids of mono-allelic genes. p0 represents the B73 allele; p1 represents the Ki11 allele.

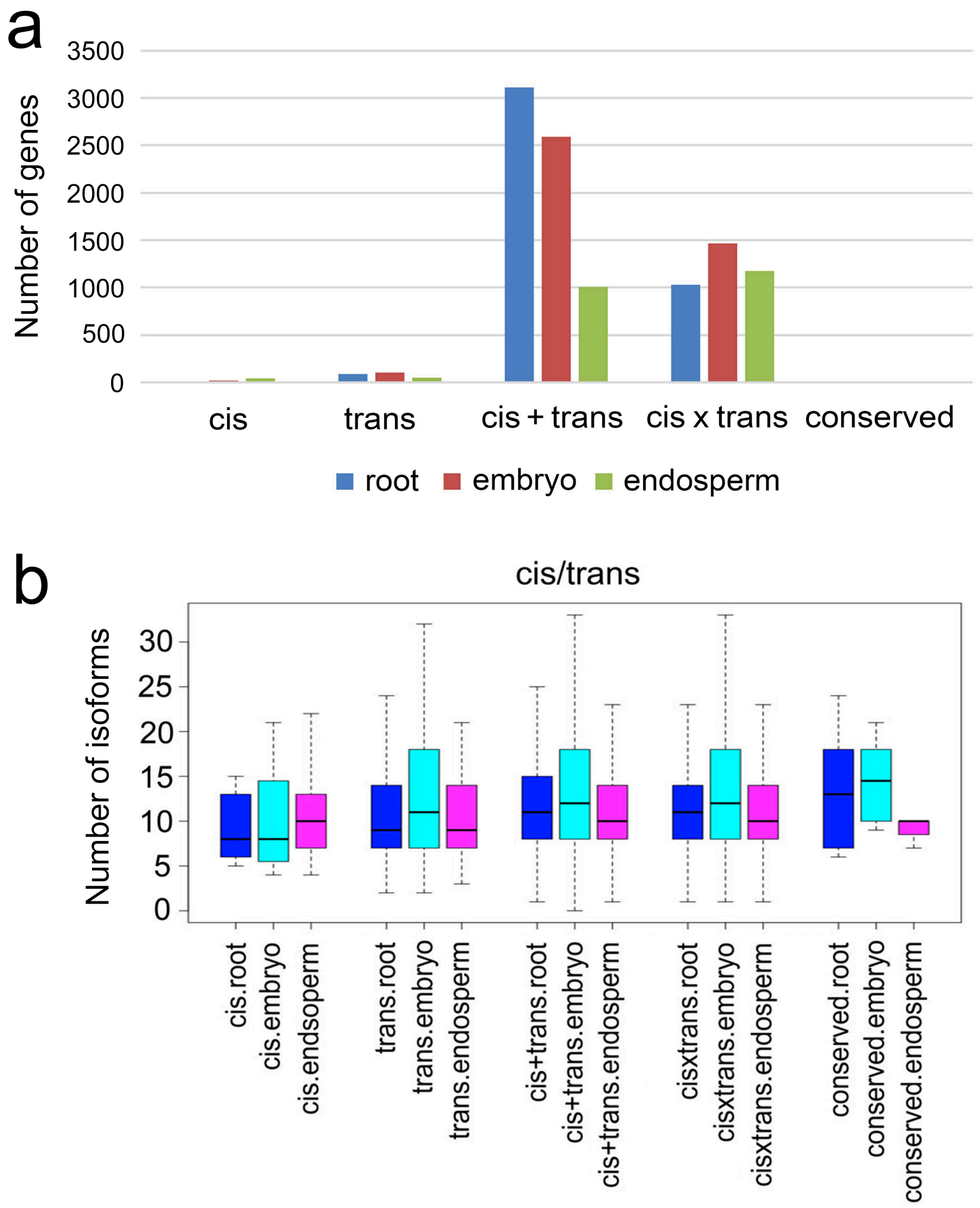


**Supplementary Figure 11. Distribution of cis-, trans-regulated genes, and number of isoforms of each category.** (a) Distribution of different categories of cis-, trans-regulated genes. (b) Number of isoforms in each category of cis-, trans-regulated genes.
